# Transient loss of Polycomb components induces an epigenetic cancer fate

**DOI:** 10.1101/2023.01.04.522799

**Authors:** V. Parreno, V. Loubière, B. Schuettengruber, M. Erokhin, B. Győrffy, M. Di Stefano, L. Fritsch, J. Moreaux, D. Chetverina, A-M. Martinez, G. Cavalli

## Abstract

Cell fate depends on genetic, epigenetic and environmental inputs that are interconnected, making it difficult to disentangle their respective contributions to cell fate decisions^1-3^, and epigenetic reprogramming is a major contributor to tumor plasticity and adaptation^4-6^. Although cancer initiation and progression are generally associated with the accumulation of somatic mutations^7,8^, substantial epigenomic alterations underlie many aspects of tumorigenesis and cancer susceptibility^9-18^, suggesting that genetic mechanisms alone may not be sufficient to drive malignant transformations^19-23^. However, whether purely non-genetic reprogramming mechanisms are sufficient to initiate tumorigenesis irrespective of mutations is unknown. Here, we show that a transient perturbation of transcriptional silencing mediated by Polycomb-Group proteins is sufficient to induce an irreversible switch to a cancer cell fate in *Drosophila*. This is linked to the irreversible derepression of genes that can drive tumorigenesis, including JNK and JAK-STAT signalling pathways and *zfh1*, the fly homolog of the ZEB1 oncogene, which we show to be a necessary driver of the cancer fate. These data show that a reversible perturbation of Polycomb-Group protein levels can induce cancer in the absence of driver mutations and suggest that this is achieved through epigenetic inheritance of altered cell fates.

## Main Text

Over the last decades, large-scale projects greatly expanded the known repertoire of cancer-associated genetic mutations affecting epigenetic factors^24-28^, including chromatin remodelers and modifiers controlling histone marks^29-31^, DNA methylation^32^, micro-RNA^33^ and 3D-genome folding^34^ that might support cancer progression^35^, corroborating the role of epigenetic aberrations in the etiology of hematological and solid malignancies^36-38^. Indeed, epigenetic modifications are used as biomarkers and are targeted by epi-drugs in cancer therapy^39^. Tumorigenesis is therefore associated with genetic as well as epigenetic determinants^40-42-43^.

The fact that several hallmarks of human cancer^43-48^ may be acquired through epigenome dysregulation suggests that epigenetic alterations play causal roles in cancer^9^. For example, pancreatic cancer metastases can arise without the emergence of driver DNA mutations, but coincide with major epigenetic changes^49,50^. Recently, epigenetic contributions were shown to play key roles in several cases of metastatic progression^51,52^. In some pediatric cancers, such as ependymomas, low numbers of mutations have been detected, suggesting that epigenetic changes may drive tumorigenesis^53^. Combined, these data led to the proposal that cancer is not a consequence of DNA mutations only^14,54^.

However, whether purely non-genetic reprogramming mechanisms are sufficient to initiate tumorigenesis irrespective of mutations remains an open question. Polycomb Group (PcG) proteins are epigenetic factors forming two main classes of complexes called Polycomb Repressive Complex 1 and 2 (PRC1 and PRC2, respectively), which are highly conserved from fly to human and play a critical role in cellular memory, by ensuring the stable repression of major developmental genes throughout development. PcG dysregulation leads to cell fate changes^55,56^, developmental transformations and is associated with multiple types of cancers^57-59^, including breast and prostate cancer^60^. PRC2 deposits the H3K27me3 repressive mark, whereas PRC1, which contains the PH, PC, PSC and the SCE subunits in flies, is responsible for H2AK118Ub deposition^61-63^. Contrasting with the extreme redundancy found in mammals^58^, most PcG components are coded by a single gene in *Drosophila*, making this system more tractable for functional studies.

### A transient epigenetic perturbation is sufficient to initiate tumors

Earlier work has shown that both null mutations or constant RNAi knock-down (KD) targeting both *ph* homologs (*ph-p* and *ph-d*) drive similar transcriptional defects and are sufficient to induce loss of differentiation and cell overproliferation^64-68^. To test whether a transient epigenetic perturbation might be sufficient to trigger an irreversible change in cell fate, we set up a thermosensitive *ph*-RNAi system allowing for the reversible KD of *ph* in the developing larval eye imaginal disc (ED, see Figure 1a, 1b; Extended Data Fig. 1). As expected, constant PH depletion induces tumors (Figure 1c,d; Extended Data Fig. 1a-d) as well as reduced adult survival (Extended Data Fig. 1e). Strikingly, a transient 24h depletion of PH at the L1 stage, where the ED starts developing, is also sufficient to trigger the formation of tumors, characterized by overgrowth, the loss of apico-basal polarity and differentiation markers (Figure 1c-e; Extended Data Fig. 1b-d). Importantly, these tumors show normal levels of the PH protein, both at day 9 (transient *ph*-KD d9) and day 11 (transient *ph*-KD d11) after egg laying (AEL) (Figure 1b, Extended Data Fig. 1a). EDs continue to grow significantly even after PH recovery (Figure 1e) and cannot differentiate (Extended Data Fig. 1d), suggesting that the tumor state is stable and maintained independently of its epigenetic trigger. Likewise, PH depletion at the L2 or early L3 stage induced tumors (Extended Data Fig. 1b-d). Of note, a 24 h exposure at 29°C at the L3 stage is sufficient to deplete PH in our system (Extended Data Fig. 1f). Consistent with the constant need of PRC1 function to prevent tumorigenesis, a transient depletion of PSC-SU(Z)2, another core subunit of PRC1 for which null mutations drive neoplastic transformation^69,70^, was also sufficient to induce tumorigenesis (Extended Data Fig. 2).

**Figure 1:**
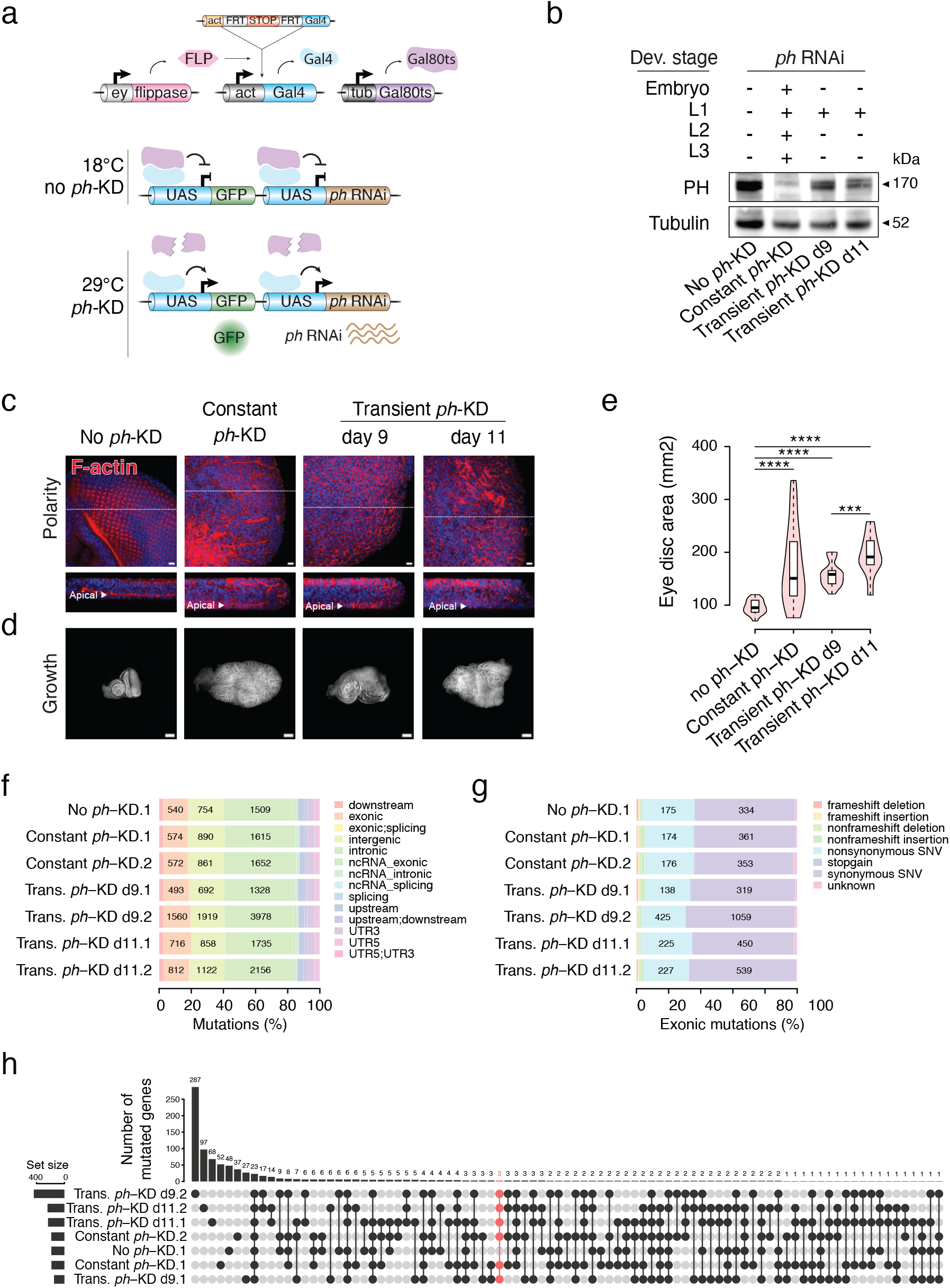
Transient PRC1 depletion is sufficient to initiate tumors. **a-**Scheme depicting the system for conditional depletion of PH. The expression of *ph-*RNAi, as well as that of the GFP marker, is under the control of UAS sequences. Cells that express *ey*-FLP (in pink) catalyze FLP-out of a transcriptional stop (in orange) in developing EDs, allowing the expression of *act*-Gal4 (in light blue). The *tub*-Gal80^ts^ (in purple) encodes a ubiquitously expressed, temperature-sensitive Gal4 repressor. At restrictive temperature (29°C), but not at 18°C, Gal80ts is inactivated. Gal4 can therefore bind to UAS sequences, allowing the expression of *ph*-RNAi and GFP that is used as a read-out of *ph*-KD. **b-** Western Blot analysis of PH expression in EDs of L3 larvae subjected to no *ph*-KD (control), constant or transient *ph*-KD at L1 stage. **c-** Confocal microscopy analysis of F-actin stained with rhodamine-phalloidin (red) showing a well-organized monolayer-stratified epithelium with apical F-actin see in planar (top) and xy cross-sections (bottom) in no *ph*-KD (control), while polarity is disrupted in constant or transient *ph*-KD EDs. DNA is stained with DAPI (blue). **d-e-** Comparative measurement of ED size quantified as an overall DAPI staining area (d) showing ED sizes (e) under no *ph*-KD (control), constant or transient *ph*-KD conditions (wilcoxon test, ***pval<1e-3, ****pval<1e-5). **f-g-** Feature distribution of all (f) or exonic (g) somatic SNPs and InDels from no *ph*-KD (control), constant or transient *ph*-KD tissues (2 biological replicates). Numbers are shown for highly represented features (>10%). **h-** Mutated gene overlaps between conditions (including frameshift, nonsynonymous and stop gain mutations). Scale bars: 10 μm (**c**), 100 μm (**d**).

To identify potential driver mutations that might arise as a consequence of PH depletion, we collected eggs from two independent batches of mated females and subjected them to a transient KD, a constant KD or no *ph*-KD (control condition), before sequencing their genomic DNA. Given the 100% penetrance of the tumor phenotype and the relatively low number of genes that can act as cancer drivers in *Drosophila*^*71*^, we reasoned that such driver mutations should be repeatedly recovered in independent tumors. However, when using the first of the two control batches as a reference in the search for somatic single nucleotide variants (SNV) or insertions and deletions (InDel) ^72^, we found that the six tumor samples showed comparable numbers of variants as well as similar features distribution compared to the second control sample (Figure 1f,g). Importantly, the large majority of variants were detected in a single sample, and few mutated genes were shared between samples (Figure 1h, see material and methods). Only three genes were found mutated in all six tumor samples and not in control, namely *spz3, CG34398* and *CG9525*, none of them having known functions in tumorigenesis. Finally, no enrichment for any specific Gene ontology (GO) was found for genes mutated in at least two *ph*-KD samples. This evidence strongly argues against the presence of recurrent driver mutations in these tumors, reminiscent of previously reports suggesting that genome instability is not a prerequisite for neoplastic epithelial growth caused by *ph*-KD^65^.

In sum, these data show that the transient depletion of tumor suppressor PRC1 components is sufficient to switch cells into a neoplastic state that is maintained even after normal levels are re-established. Since the same genotype can generate both a normal phenotype or a tumor depending on a transient gene regulatory modification, we defined these tumors as epigenetically-initiated cancers (EICs).

### EICs are characterized by activated JAK-STAT and JNK signaling

To tackle the molecular mechanisms leading to the formation of EICs, we compared the transcriptomes of control (no *ph*-KD), transient and constant *ph*-KD to temperature-matched controls, generated with a similar RNAi system targeting the *white* gene, which is dispensable for normal eye development (see material and methods, Figure 2, Extended Data Table 1). As expected, both systems are hardly distinguishable at 18°C (no RNAi), as well as upon transient *w*-KD (Figure 2a; Extended Data Fig. 3a). Consistent with our previous work^64,68^, constant *ph*-KD is associated with the up-regulation of 340 genes – including canonical PcG targets such as Hox and developmental Transcription Factor (TF) genes – and the down-regulation of 2,110 genes, including most key regulators of ED development (Figure 2a; Extended Data Table 1; Extended Data Fig. 3b). Importantly, only a subset of these genes were also differentially expressed in transient *ph*-KD at d9 AEL (256 and 812, respectively), and even less at later d11 AEL (154 and 446, respectively), suggesting a progressive yet incomplete rescue of the transcriptome (Figure 2a; Extended Data Fig. 3, 4a; Extended Data Table 1). Therefore, a substantial portion of the transcriptome can be restored upon reinstating normal levels of PH.

**Figure 2:**
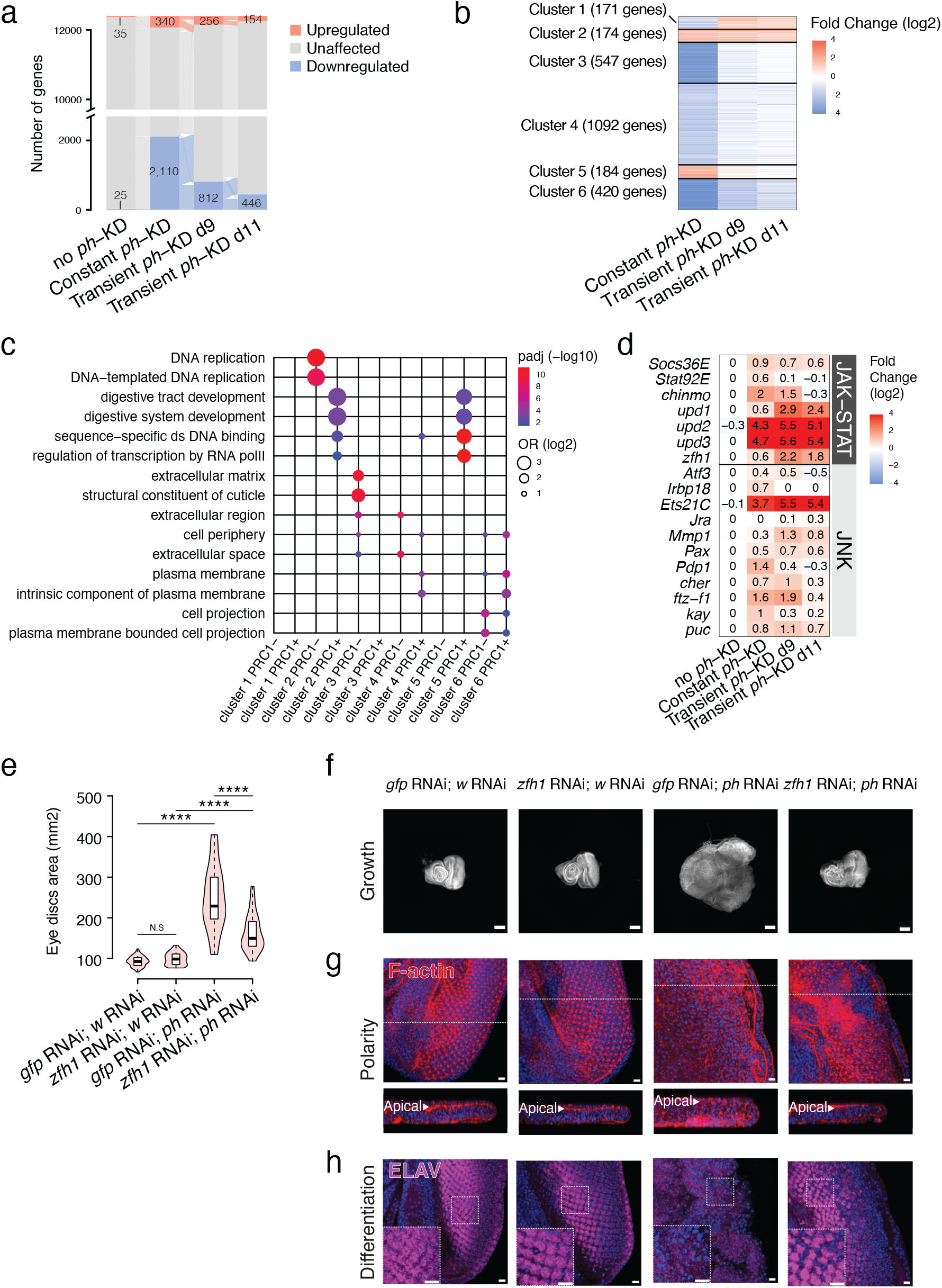
EICs maintain a subset of the dysregulated tumoral transcriptome and their maintenance requires the ZFH1 TF. **a-** Alluvial plot displaying the dynamics of transcriptional states between no *ph*-KD (control), constant and transient *ph*-KD. Transitions between upregulated (orange), unaffected (grey) and downregulated (blue) states are indicated by thin lines of the same respective colors. **b-** Clustering of genes differentially expressed in constant and/or transient *ph*-KD, but not in the control no *ph*-KD. **c-** Gene ontologies significantly associated with each gene cluster, further stratified for the presence or absence of PH peaks located in the gene body or up to 2.5 kb upstreat to the TSS. **d-** Transcriptional fold changes in genes involved in JAK-STAT (top) and JNK (bottom) signaling pathways upon PH depletion. **e-** Comparative size of *ph*-RNAi EDs in the presence or absence of *zfh1*-RNAi and the corresponding control RNAi (targeting the *white* and *gfp genes*, respectively). **f-h** stainings of *ph*-RNAi EDs in the presence or absence of *zfh1*-RNAi and the corresponding control RNAis (directed against *white* and *gfp*, respectively), marking the DAPI area (growth, **f**), F-actin (polarity, **g**) and neuronal marker ELAV (differentiation, **h**). Scale bars: 100 μm (**f**), 10 μm (**g**,**h**).

Hierarchical clustering of differentially expressed genes identified six clusters with specific dynamics (Figure 2b; Extended Data Table 2). Genes of Cluster 3, 4 and 6 are downregulated and gradually or fully return to normal levels of expression (Figure 2b; Extended Data Table 2). In contrast, upregulated genes (Cluster 1, 2 and 5) show 3 different patterns (Figure 2b; Extended Data Table 2). To identify direct PcG target genes correlated to these patterns, we performed PH ChIP-Seq, and H3K27me3 and H2AK118Ub CUT&RUNs – the canonical marks of PcG-mediated transcriptional repression – in control EDs (see material and methods, Extended data 5a and Extended Data Table 2). Cluster 1 genes are specifically upregulated in EICs, are not enriched for any specific gene ontology (GO) and are generally not targeted by PcG (Figure 2b,c and Extended Data Table 2), suggesting that their up-regulation is mostly an indirect consequence of PH depletion. In contrast, direct PcG target genes are over-represented in clusters 2 and 5 (Extended Data Fig. 5b), which are enriched for GOs related to developmental TFs (Figure 2b, c).

Cluster 5 includes many TFs that are bound by both PRC1 and PRC2 and correspond to canonical PcG targets such as *en, eve, wg, Scr, Su(z)12, Ubx* and *wg*. Their progressive recovery of control levels of expression at d9 or d11 AEL precludes these genes from being required for the maintenance of EICs. On the other hand, cluster 2 contains genes which share the interesting feature of being irreversibly upregulated, irrespective of PH recovery. Among them, we noted an over-representation of direct PcG target genes involved in JAK/STAT (*upd1, upd2, upd3*, s*ocs36E, zfh1, chinmo*) and JNK (*Ets21C, puc*) signaling pathways (Figure 2d, Extended Data Table 2), which were shown to play an active role in the development of tumors in various *Drosophila* cancer models^66,69,73^. In parallel, genes from cluster 2 which are not bound by PRC1 are enriched for DNA-replication related GOs (Figure 2c), and likely are induced by indirect mechanisms. Cluster 2 also contains the Insulin-like peptide 8 (*Ilp8*) gene responsible for developmental delay of tumor-bearing larvae^63^, whose irreversible induction is a strong signature of abnormal JNK signaling activation (Extended Data Tables 1 and 2).

Irreversibly upregulated genes of cluster 2 contain several key drivers of tumorigenesis in *Drosophila* conserved in mammals. Among these, we noticed the presence of *zhf1*, a TF that is both a direct PcG target (Extended Data Fig. 5a), is activated by multiple signaling pathways, including JAK/STAT and Notch signaling and which can sustain self-renewal and tumor growth^74-76^. Furthermore, its mammalian homolog ZEB1 can induce epithelial tomesenchymal transition^77^. Consistent with its transcriptional upregulation, ZFH1 protein is increased in both constant and transient tumors (Extended Data Fig. 4b,c). Importantly, *zfh1* RNAi reduces the growth of *ph*-dependent tumors and partially restores cell polarity and photoreceptor differentiation (Figure 2e-h), indicating that it is a *bona fide* driver of the tumor phenotype. Together, these results demonstrate that EICs are driven by a restricted set of irreversibly upregulated genes, including JNK and JAK/STAT signaling components, rather than by the vast pleiotropic dysregulation of cancer genes that is observed upon constant PH depletion. Furthermore, we sought to investigate why this subset of genes remains irreversibly upregulated after restoration of normal PH levels.

### Irreversible transcriptional changes drive *ph*-dependent tumorigenesis

To identify the features that discriminate reversible genes from irreversible ones, we focused on cluster 2 (irreversible) and cluster 5 (reversible) genes that were either bound by PH or enriched for H3K27me3 or H2AK118Ub (the canonical marks of PcG-mediated transcriptional repression, Extended Data Table 2). As expected, reversible genes (n= 68) show no expression changes after transient *ph*-KD, whereas irreversible genes (n= 61) remain upregulated (Extended Data Table 2, Figure 3a). PRC1-target genes are transcribed at similarly low levels in both clusters in the absence of *ph*-KD, and induced at comparable levels upon constant *ph*-KD; ruling out that higher transcriptional levels are the reason for irreversible genes being unable to recover normal transcription after transient *ph*-KD (Figure 3a,b).

**Figure 3:**
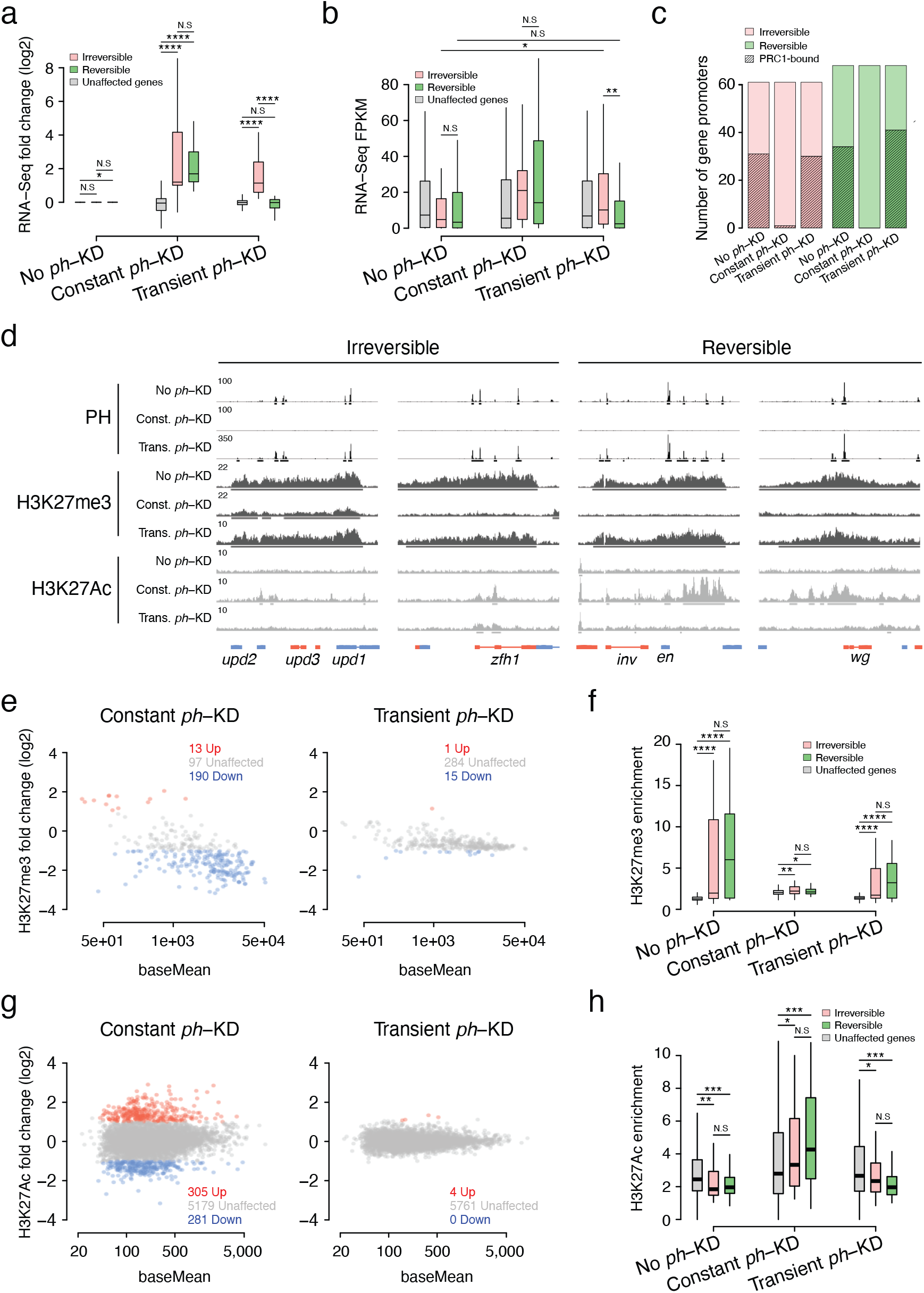
Stable transcriptional activation of irreversible genes despite the presence of Polycomb chromatin features. **a-, b-** Fold Changes (**a**) and FPKMs (**b**) of irreversible (pink) and reversible (green) genes. All unaffected genes are used as reference (wilcoxon test, N.S-not significant, *pval<5e-2, **pval<1e-2, ***pval<1e-3, ****pval<1e-5). **c-** Number of irreversible (pink) and reversible (green) gene promoters that are bound by PRC1 (black shading lines) in no *ph*-KD (control), constant or transient *ph*-KD. **d-** Screenshot of ChIP-Seq profiles for PH, H3K27me3 and H3K27Ac CUT&RUNs at representative genes activated irreversibly (left) or reversibly (right) under the indicated conditions. **e-** Differential analysis of H3K27me3 domains that are unaffected (grey) or exhibit decreased (blue) or increased (orange) enrichment upon constant (left) or transient (right) PH depletion. **f-** Quantification of H3K27me3 at irreversible (pink) and reversible (green) gene bodies, using all unaffected genes as reference (grey). Wilcoxon test, N.S-not significant, *pval<5e-2, **pval<1e-2, ***pval<1e-3, ****pval<1e-5. **g-** Differential analysis of H3K27Ac peaks that are unaffected (grey) or exhibit decreased (blue) or increased (orange) enrichment upon constant (left) or transient (right) PH depletion. **h-** Quantification of H3K27Ac at irreversible (pink) and reversible (green) gene promoters, using all unaffected genes as reference (grey). Wilcoxon test, N.S-not significant, *pval<5e-2, **pval<1e-2, ***pval<1e-3, ****pval<1e-5.

We then explored the possibility that chromatin might not be correctly re-established at irreversible genes. We performed ChIP-Seq and CUT&RUN in constant and transient (d11) *ph*-KD EICs. PH ChIP-Seq showed a loss of PH binding for both reversible and irreversible gene sets upon constant *ph*-KD, as expected, while PH binding was remarkably recovered after transient *ph*-KD (Figure 3c, d; Extended Data Fig. 5h). CUT&RUNs against H3K27me3 indicates a stark decrease upon constant *ph*-KD and a reciprocal increase in H3K27Ac – its activating counterpart – both at reversible and irreversible genes (Figure 3 d-h and Extended Data Fig. 5c) and a similar yet weaker trend was observed for H2AK118Ub (Extended Data Fig. 5e-f). Interestingly, both clusters show comparable H3K27me3 levels after transient *ph*-KD, with a concomitant decrease in H3K27ac (Figure 3d,g,h). Inspection of individual loci showed that qualitative and quantitative recovery is generally observed at both reversible and irreversible genes (Figure 3d-h and Extended Data Fig. 5c,e,f,h). Taken together, these results indicate a suprising uncoupling between the impact of transient *ph*-KD on transcription and chromatin, whereby irreversible transcriptional changes drive tumorigenesis despite the re-establishment of an essentially normal chromatin landscape.

### Transcriptional repressor binding distinguishes reversible and irreversible genes

To test whether differential levels of PRC1 binding might explain the repression of reversible genes upon transient *ph*-KD, we focused on reversible (n= 32) and irreversible (n= 26) genes whose promoters are bound by PH (see material and methods, Extended Data Table 2). Despite showing similar transcriptional levels (Figure 4a), PH ChIP-Seq in control EDs (no *ph*-KD) indicates that PH binding and H2AK118Ub levels are significantly stronger at the TSSs of reversible genes compared to irreversible ones (Figure 4b, Extended Data 5g). A similar trend is observed for H3K27me3, although not significant (Extended Data 5d). This was futher confirmed using available ChIP-Seq data from WT EDs^64,68^ for the two other tumor-suppressor PRC1 members, PC and PSC, and the PRC2 member SU(Z)12, suggesting that the fully assembled PRC1 binds together with PRC2 to these promoter regions (Figure 4c,d). We therefore analyzed the binding profiles of sequence-specific DNA binding factors that are known to recruit PcG proteins – namely Pleiohomeotic (PHO), Sp1-like factor for pairing sensitive-silencing (SPPS)^78^, Combgap (CG)^79^ and Trithorax-like (TRL)^80^ – and found that all 4 show similar trends, with increased binding at the promoters of reversible genes (Figure 4e).

**Figure 4:**
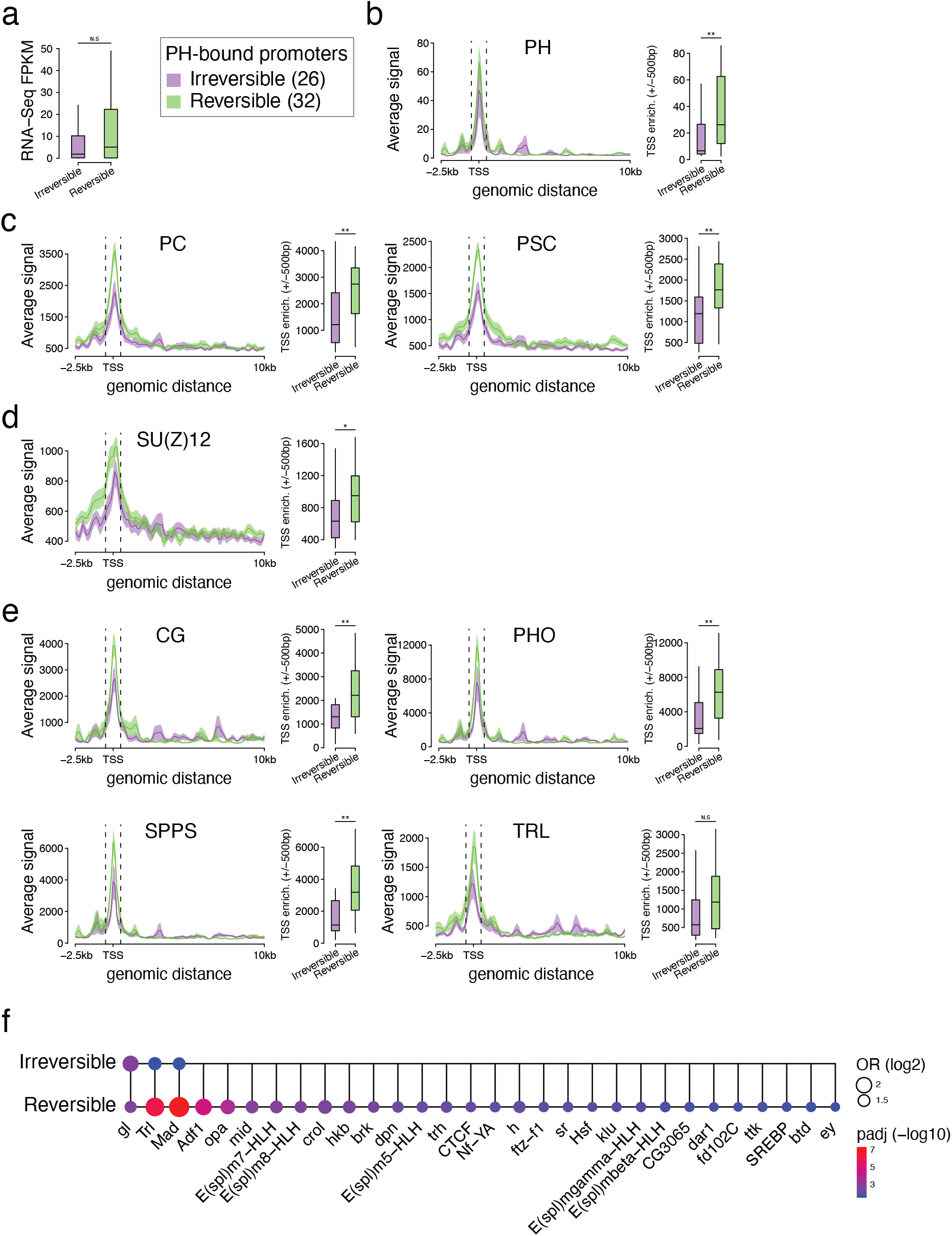
Irreversible genes are identified by lower levels of binding of Polycomb components and PcG recruiters. **a-** Boxplots of RNA levels in FPKMs of irreversible (purple) and reversible (green) PH-bound genes (wilcoxon test, N.S-not significant). **b-e-** Metaplots of average CHIP-Seqs for PRC1 (PH, PC, PSC) and PRC2 (SU(Z)12) subunits, as well as PcG recruiter proteins (CG, PHO, SPPS, TRL) to irreversible (purple) and reversible (green) genes. Quantifications are shown on the right (wilcoxon test, N.S-not significant, *pval<5e-2, **pval<1e-2, ***pval<1e-3, ****pval<1e-5). **f-** Comparative enrichment of DNA consensus binding motifs at irreversible and reversible gene promoters (−200bp/+50bp TSSs), irrespective of PH binding. All unaffected gene promoters were used as a control set.

Finally, DNA sequence analysis shows that consensus motifs for the TRL, and ADF^81^ recruiters of PcG components are enriched at reversible genes (Figure 4f)^82,83^. TSSs of reversible genes are significantly enriched for consensus of several transcriptional repressors, such as E(SPL)bHLH repressors including hairy (*h*), as well as HKB, BRK, DPN, KLU, TTK, DAR1, KLF15 and GRH (Figure 4d).

Together, these results indicate a tight equilibrium, wereby less potent recruitment of PRC1 coupled with reduced binding of independent transcriptional repressors make irreversible genes more sensitive to transient *ph*-KD, while the upregulation of reversible genes requires constant *ph*-KD.

### EICs are autonomous, immortal tumors that become more aggressive over time

Most EICs-bearing larvae die after d11 AEL, preventing the study of tumor development over time. To circumvent this limitation, allografts of imaginal disc tissue into the abdomen of adult *Drosophila* hosts are commonly used to assess the tumorigenic potential of a tissue, and we previously showed that *ph* mutant EDs continuously grow until they eventually kill the host^66^. To be able to track transplanted EICs, we developed a variant of our themosensitive system that constitutively expresses GFP in the eye, while an UAS-RFP cassette can be used as a reporter of ongoing *ph*-KD (Extended Data Fig. 6a, f-i). Importantly, this system induces EICs with similar penetrance, morphological and transcriptional defects compared to the previous one (Extended Data Fig. 6b-e; Extended Data Table 3). We then performed a series of allografts using this line (Figure 5a). Of note, host flies were always kept at a restrictive temperature after transplant (18°C), precluding that further overgrowth would result from ongoing *ph*-RNAi.

**Figure 5:**
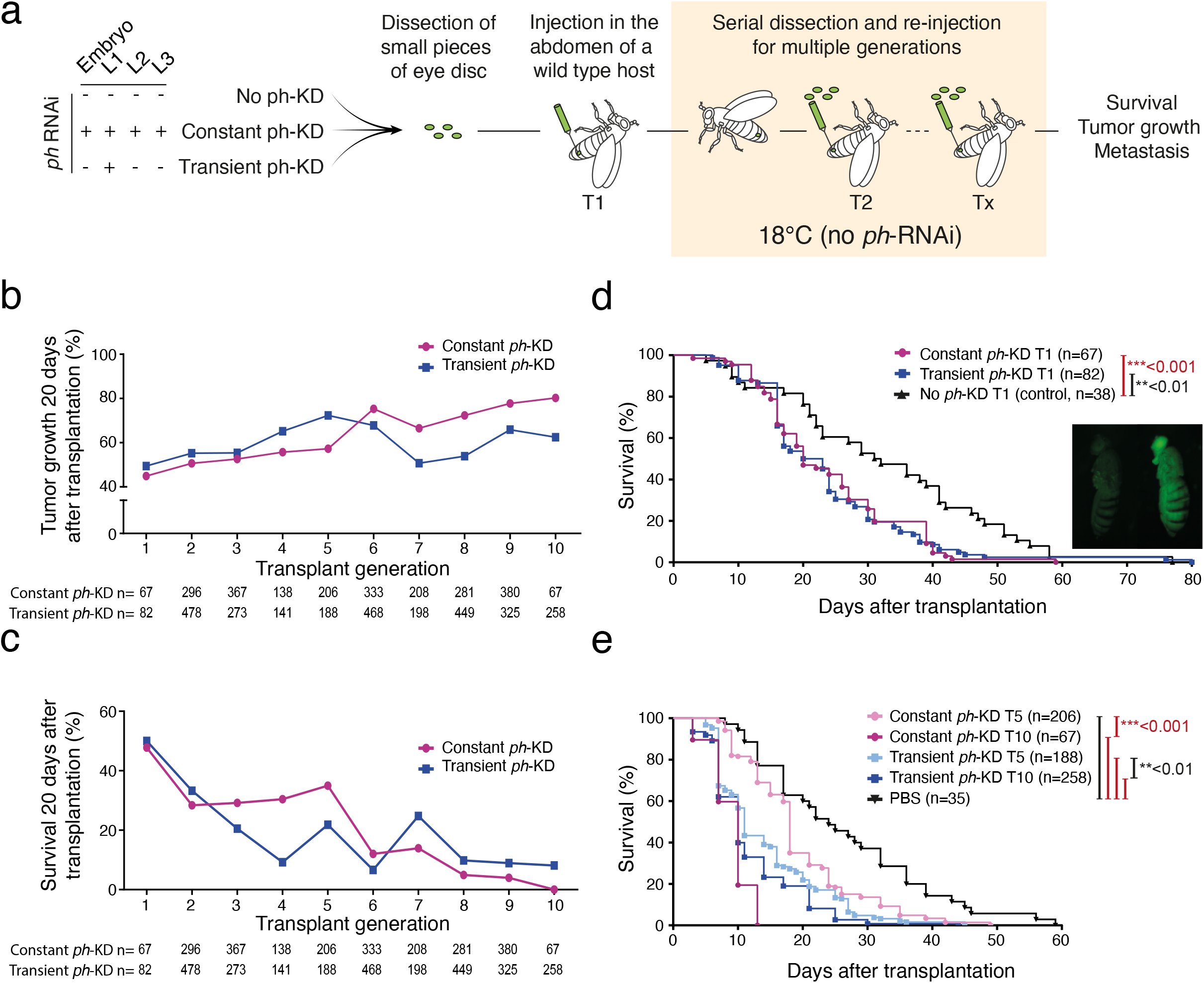
EICs generate immortal malignancies. **a-** Schematic overview of experimental allograft workflow. GFP positive Flies of the same genotype are subjected to different temperature conditions at L1 stage, namely 18°C as control, constant PH depletion at 29°C during larval stages, or transient, 24H PH depletion at 29°C. EDs are dissected from donor larvae and allografted repeatedly into the abdomen of host flies until T10 (≈ 3 months). All allograft experiments are performed at 18°C to avoid *ph*-RNAi expression. **b-, c-** Tumor growth measured as a percentage of flies showing tumoral progression 20 days after transplantation (**b**) and survival of host flies 20 days after allograft of constant (purple) or transient (blue) *ph*-KD tumors during 10 rounds of transplantation (**c**). **d-, e-** Cumulative survival rate as percentage of flies calculated for 80 days after the first transplantation (**d**) and comparative cumulative percentage of flies that develop tumors calculated for 60 days after the fifth and the tenth transplantations of constant (purple and pink respectively) or transient (pale and dark blue respectively) *ph*-KD transplants. Significant differences were assessed using Log-rank test. n indicates the number of host flies analyzed.

Constant *ph*-KD tumors were able to expand and invade a fraction of injected host flies at restrictive temperature (Figure 5b). Transient *ph*-KD EICs behaved similarly, indicating that their overgrowth results from an autonomous, stably acquired state (Figure 5b). To quantitatively measure tumor growth over time, we set up an arborescent allograft scheme, allowing us to trace the tumor of origin (Extended Data Fig. 7a). Tumors derived from both, constant or transient PH depletion, maintained their ability to expand in host flies over 10 rounds of transplantation (about 3 months). Tumor growth penetrance – defined as the percentage of hosts flies bearing GFP positive cells 20 days after transplantation – did not decrease but rather increased over generations of transplantation (Figure 5b), as the survival of host flies kept on decreasing (Figure 5c). Median lifespan of the hosts after the first transplant with constant or transient EICs was of 20 and 21,5 days, respectively (Figure 5d), compared to 31,5 days for control tissues. In contrast, hosts transplanted with the fifth generation of allografts survived 18 (constant *ph*-KD) and 11 days (transient *ph*-KD, d11), and only 10 days for tenth generation allografts (Figure 5e). Furthermore, tumors produced metastases in regions and organs located far away from the injection site, with an increasing penetrance (Extended Data Fig. 7b, c).

These results suggest that, rather than being a milder form of neoplastic transformation, EICs tumorigenic potential is autonomously maintained, increases over time, and can propagate months after *ph*-RNAi has been removed.

## Discussion

Despite great interest in epigenetic determinants of disease states, it is difficult to discriminate between genetic, environmental and cell-intrinsic epigenetic contributions, which might cooperate to induce tumorigenesis^52^. Our model system shows that upon transient depletion of PRC1 subunits, cells derail from their normal fate and undergo neoplastic transformation (Extended Data Fig. 8). This transformation coincides with the irreversible activation of genes including key JNK and JAK/STAT pathway members, which in turn might sustain cell growth, proliferation, loss of cell polarity, cell migration and cytokine activity even in the presence of normal PRC1 function. One main difference between these irreversibly activated genes and reversible PcG target genes is a more potent PcG binding on the latter ones, consistent with higher levels of PcG recruiters. We posit that, even if PRC1 is wiped out from these genes upon depletion, stronger PcG binding, perhaps concomitant with the function of additional repressive TFs, facilitates the restoration of a repressive chromatin environment once PRC1 levels are re-established. In contrast, weaker PRC1 binding at irreversible genes might not suffice to fully restore silencing, resulting in a self-sustaining aberrant cell state that stimulates tumor progression (Extended Data Fig. 8). Of note, even if some of these genes are rather strongly transcribed, the levels of H3K27Ac on their promoters and gene bodies are distinctly lower compared to a random set of equally transcribed genes (not shown). It will be interesting to analyze the basis for this difference and its significance.

Although the system described here has not been analyzed before, previous work has shown that self-sustaining alternate cell states can be triggered by transient perturbations in immortalized breast cells^84^ or other cultured cells^85,86^, including neural progenitor cells subjected to transient inhibition of the PRC2 complex^87^. PRC2 impairment in mouse striatal neurons induces progressive neurodegeneration by triggering a self-sustaining transcription derailment program over time^88^. Furthermore, KO or transient chemical inhibition of PRC2 also led cells to enter a quasi mesenchymal state that depends on ZEB1, the mouse homolog of fly *Zfh1*, is highly metastatic and associates with poor patient survival^77^. Therefore, epigenetic events might play a major role at early stages of oncogenesis or during tumour progression in some mammalian cancers^54,89,90^. Our survey of a large database of different types of solid cancers (Extended Data Fig. 9) as well as of data from several Multiple Myeloma patient cohorts (Extended Data Fig. 10) indicates that low expression levels of genes encoding canonical PRC1 subunits is associated with several tumours and with poor patient prognosis, consistent with a putative suppressive role for PRC1 in some tumor types. Future work might address the role of epigenetic perturbations in these tumors and in other physiological processes.

## Supporting information

Ext Data Table 1

Ext Data Table 2

Ext Data Table 3

## Methods

### Fly strains

Fly lines were obtained from Bloomington Drosophila Stock Center (BL) and Vienna BioCenter Core Facilities (VDRC). All genotypes are indicated using *Drosophila* genetics annotation, i.e. a “;” indicates a different chromosome. Genotypes are indicated starting from items located in the sex chromosomes on the left, items on autosome 2 in the center and on autosome 3 on the right. A blank space is left when no specific items are present in one chromosome. For heterozygous conditions, each allele is indicated and a “/” separates the two alleles on homologous chromosomes.

The following fly strains were used for immunostaining, RNA-seq and CUT&RUN experiment:

y[1] w[*] P{ry[+t7.2]=ey-FLP.N}2 P{w[+mC]=GAL4-Act5C(FRT.CD2).P}D ; ; M{w[+mC]=UAS-NowGFP-NLS}ZH-86Fb (BL#64095, driver line expressing Gal4 in the developing eye disc. Expresses GFP under the control of Gal4).

; P{w[+mC]=tubP-GAL80[ts]}20 ; UAS-phRNAi (derived from BL#7019 and VDRC#50028; used for *ph*-KD).

; P{w[+mC]=tubP-GAL80[ts]}20 ; UAS-wRNAi (derived from BL#7019 and BL#33623, used for *white*-KD and the generation of temperature-matched control transcriptomes).

; UAS psc-su(z)2 RNAi ; P{w[+mC]=tubP-GAL80[ts]}ncd[GAL80ts-7] (derived from BL#38261, VDRC#100096 and BL#7018, used for *Psc/Su(z)2*-KD).

The following fly lines were used for the *Zfh1* rescue experiment:

y[1] w[*] P{ry[+t7.2]=ey-FLP.N}2 P{w[+mC]=GAL4-Act5C(FRT.CD2).P}D ; ; M{w[+mC]=UAS-NowGFP-NLS}ZH-86Fb (BL#64095, driver line expressing Gal4 in the developing eye disc. Expresses GFP under the control of Gal4).

; UAS zfh1 RNAi (VDRC#103205, used for *zfh1*-KD).

; UAS GFP RNAi (BL#9391, control RNAi).

; ; UAS-wRNAi (BL#33623, control RNAi).

; ; UAS-phRNAi (VDRC#50028, used for *ph*-KD).

The following lines were used for the allograft assay and gDNA sequencing:

eyFLP, UbiP63E(FRT.STOP)Stinger ; ; Act5C-Gal4(FRT.CD2), UAS-RFP /TM6BTb (derived from BL#5580, BL#32249 and BL#30558; driver line expressing Gal4 and GFP in the developing eye disc. Expresses RFP under the control of Gal4).

; P{w[+mC]=tubP-GAL80[ts]}20 ; UAS-phRNAi (derived from BL#7019 and VDRC#50028; used for *ph*-KD). w[*] ; P{w[+mC]=His2Av-mRFP1}III.1 (BL#23650).

### Experimental crosses

Fly lines were maintained and amplified at 25°C on standard food medium. For all crosses, egg laying was performed for 4h at 18°C, before applying the following temperature shifts:

- No *ph*-KD: 18°C, dissection 10 days After Egg Laying (AEL).
- Constant *ph* -KD: 29°C, dissection 4 days AEL.
- Larval depletion: 48 hours at 18°C, 29°C until dissection 5 days AEL.
- Transient *ph* -KD: 18°C for 48h, 29°C for 24h, 18°C until dissection at day 9 (d9) or day 11 (d11) AEL.
- Transient *ph* -KD L2 stage: 18°C for 96 hours, 29°C for 24 hours, 18°C until dissection at day 8 AEL.
- Transient *ph* -KD L3 early stage: 18°C for 120 hours, 29°C for 24 hours, 18°C until dissection 8 days AEL.
- Transient *ph* -KD L3 late stage: 18°C for 168 hours, 29°C for 24 hours, 18°C until dissection 8 days AEL.
- Transient *Psc*-KD: 18°C for 48 hours, 29°C for 48 hours, 18°C until dissection 8 days AEL.

### Immunostaining procedures

Eye-antennal imaginal discs from third instar female larvae were dissected at RT in 1x PBS and fixed in 4% Formaldehyde for 20 min. Tissues were permeabilized for 1h in 1x PBS + 0.5% Triton X-100 on a rotating wheel. Permeabilized tissues were blocked 1 hour in 3% BSA PBTr (1x PBS + 0.1% Triton X-100), and incubated O/N on a rotating wheel at 4°C with primary antibodies diluted in PBTr + 1% BSA. The following primary antibodies were used: goat anti-PH ^1^ (1:500), mouse anti-ELAV (1:1000, DSHB, 9F8A9), mouse anti-ABD-B (1:1000, DSHS, 1A2E9), chicken anti-GFP (1:500, Life Technologies, D27C4), rabbit anti-zfh1^2^ (1:2000). Then, sample were washed in PBTr before adding secondary antibodies or rhodamine phalloidin in PBTr and 2 hours at RT on rotating wheel. The following secondary antibodies were used: donkey anti-goat Alexa Fluor 555 (1:1000, Invitrogen, A-21432), donkey anti-mouse Alexa Fluor 647 (1:1000, Invitrogen, A-31571), donkey anti-chicken (1:1000, Clinisciences, 703-546-155), donkey anti-rabbit Alexa Fluor 555 (1:1000, Invitrogen, A-31572). F-actin was stained by adding rhodamine phalloidin Alexa Fluor 555 (1:1000, Invitrogen, R415) or Alexa Fluor 488 (1:1000, Invitrogen, A12379). Tissues were washed in PBTr. DAPI staining were done at a final concentration of 1 µg/mL during 15 minutes. Discs were washed in PBTr. Discs were mounted in Vectashield medium (eurobio scientific, catalog no. H-1000-10) and microscopy acquisition was performed on Leica SP8-UV confocal microscope. Eye disc area was measured by defining the limits of the tissue using Fiji on at least 30 discs.

### RNA-seq

Third instar female larvae were dissected in Schneider medium on ice. Total RNA was extracted using TRIzol reagent. RNA purification was performed using RNA Clean & Concentrator kit (Zymo Research, #R1015). Finally, poly-A RNAs selection, library preparation and illumina sequencing (20M paired-end reads, 150nt) were performed by Novogene (https://en.novogene.com/). All experiments were performed in triplicate.

### Genomic DNA sequencing

Genomic DNA was isolated using QIAamp DNA Micro Kit (Qiagen) following manufacturer’s instructions. For each biological replicate around 70 eye discs from female wandering larvae were dissected. Genomic DNAs were processed for library preparation by Novogene (https://en.novogene.com/). Briefly, genomic DNA was fragmented to an average size of ∼350bp and then processed for DNA library preparation by following manufacturer (Illumina) paired-end protocols. Sequencing was performed using Illumina Novaseq 6000 platform to generate 150bp paired-ends reads with a coverage of at least 10x for 99% of the genome.

### Western blot

About 150 eye-antennal imaginal discs were dissected in Schneider medium on ice per replicate. In order to collect enough material, eye-antennal discs were dissected in batches, snap frozen in liquid nitrogen and stored at -80°C. Discs were homogenized with a Tenbroeck directly in RIPA lysis buffer (50 mM Tris pH 7.5, 150 mM NaCl, 1% NP40, 0.5% Na-deoxycholate, 0.1% SDS, 2X protease inhibitor) and incubated on ice for 10min. If necessary, a second round of mechanical dissociation was performed. Samples were centrifuged 10 min at 10,000 x g at 4°C and supernatant was transferred to a fresh tube. Proteins were quantified using BCA protein assay and 10 µg were used per gel lane, before 20 min of migration at 200V in MES 20X migration buffer and 1h transfer (1A). Membranes were blocked for 1 hour in PBS + 0.2% Tween + 10% milk powder at RT, incubated O/N with primary antibodies in PBS + 0.2% tween at 4°C on shaker and washed in PBS+0.2% Tween. The following primary antibodies were used: rabbit anti-PH (1:200), rabbit anti-zfh1 ^2^ (1:2000), mouse anti-beta tubulin (1:5000, DSHB, AA12.1). HRP-conjugated secondary antibodies were incubated with the membrane for 2 hours at RT. The following secondary antibodies were used: goat anti-rabbit (1:15 000, Sigma, A0545), Rabbit anti-mouse (1:15 000, Sigma A9044). Membranes were washed in PBS+0.2% Tween and revealed using Super Signal West Dura kit (Pierce) and Chemidoc Bio-rad.

### Allograft assay

Allografts were performed as previously described (Rossi and Gonzalez 2015). In brief, EDs were dissected in PBS from 3^rd^ instar female larvae, cut into pieces and injected into the abdomen of adult female hosts genotype (BL#23650). Flies were monitored every two days and tumors were dissected and re-injected when the host abdomen was fully GFP.

### ChIP-seq experiments

ChIP-Seq on 3^rd^ instar imaginal eye discs were essentially performed as described earlier ^3^, with minor modifications. 400 eye discs were used per replicate. When necessary, several batches of dissection/collection were snap frozen in liquid nitrogen and stored at −80°C to collect enough material. Chromatin was sonicated using a Bioruptor Pico (Diagenode) for 10 min (30sec in, 30sec off). PH antibodies were diluted 1:100 for IP. After decrosslinking, DNA was purified using MicroChIP DiaPure columns from Diagenode. DNA libraries for sequencing were prepared using the NEBNext® Ultra™ II DNA Library Prep Kit for Illumina. Sequencing (paired-end sequencing 150bp, approx. 4Gb/sample) was performed by Novogene (https://en.novogene.com/).

### CUT&RUN experiments

CUT&RUN experiments were performed as described by Kami Ahmad in protocols.io (https://dx.doi.org/10.17504/protocols.io.umfeu3n) with minor modifications. 50 eye discs were dissected in Schneider medium, centrifuged for 3 min at 700g and washed twice with wash+ buffer before addition of Concanavalin A-coated beads. MNase digestion (pAG-MNase Enzyme from Cell Signaling) was performed for 30 min on ice. After ProteinaseK digestion, DNA was recovered using SPRIselect beads and eluted in 50ul TE. DNA libraries for sequencing were prepared using the NEBNext® Ultra™ II DNA Library Prep Kit for Illumina. Sequencing (paired-end sequencing 150bp, approx. 2Gb/sample) was performed by Novogene (https://en.novogene.com/).

### Bioinformatic analyses

All in-house bioinformatic analyses were performed in R version 3.6.3 (URL https://www.R-project.org/). Computations on genomic coordinate files and downstream computations were conducted using the data.table R package (data.table: Extension of ‘data.frame’. https://r-datatable.com, https://Rdatatable.gitlab.io/data.table, https://github.com/Rdatatable/data.table, v1.14.2).

### gDNA processing and mapping of somatic variants

gDNA variants calling was performed by Novogene (https://en.novogene.com/). Briefly, base calling was performed using Illumina pipeline CASAVA v1.8.2, and subjected to quality control uqing fastp with the following parameters: -g -q 5 -u 50 -n 15 -l 150 --min_trim_length 10 --overlap_diff_limit 1--overlap_diff_percent_limit 10. Then, sequencing reads were aligned to the dm6 version of the *Drosophila* genome using Burrows-Wheeler Aligner (BWA) with default parameters and duplicate reads were removed using samtools and PICARD (http://picard.sourceforge.net). Raw SNP/InDel sets were called using GATK with the following parameters: --gcpHMM 10 -stand_emit_conf 10 -stand_call_conf 30. Then, SNPs were filtered using the following criteria: SNP: QD<2, FS>60, MQ<30, HaplotypeScore>13, MappingQualityRankSum<-12.5, ReadPosRankSum<-8; For INDEL variants, following criteria were used: QD<2, FS>200, ReadPosRankSum<-20. UCSC known genes were used for gene and region annotations. Finally, variants were compared to the first batch of control, no *ph*-KD ED in the search for bona fide de novo mutations using the MuTect2 module of the GATK package. Only SNP and INDEL variants that passed Mutect2 filtering (FILTER= “PASS”) were considered for downstream analyses.

### RNA-seq processing and differential analysis

After initial quality checks of newly generated data using fastqc (http://www.bioinformatics.babraham.ac.uk/projects/fastqc/), paired-end reads were aligned to a custom index consisting of the dm6 version of the *Drosophila* genome together with GFP, EGFP and mRFP1 sequences, using the align function from the Rsubread R package ^4^ (v2.0.1) with the following parameters: maxMismatches = 6, unique = TRUE. Then, aligned reads were counted for each *D. melanogaster* transcript (dmel_r6.36 annotation) using the featureCounts function from the Rsubread R package (v2.0.1, isPairedEnd = TRUE) and differential expression analysis was performed using the DESeq2 R package ^5^ (v1.26.0, design = ∼replicate+condition). For each condition, *ph*-RNAi were compared to temperature-matched *w*-RNAi controls.

Then, genes that were not affected at 18°C (padj>=0.05) but differentially expressed in at least one of the other *ph*-RNAi conditions (padj<0.05 and |log2Fold2FoldChange| >1) were considered for clustering. Log2FoldChange values were clipped at 5^th^ and 95^th^ percentile, and clustered using the supersom function from the kohonen R package ^6^ (v3.0.10). Given that D9 and D11 transient *ph*-KD yielded substantially similar transcriptomes, a two-layer self-organizing map was trained (layer 1: Constant *ph*-KD; layer2: D9 and D11 transient *ph*-KD) with similar weights for the 2 layers, using a 3×2 grid (topology = hexagonal, toroidal = TRUE).

### CUT&RUN and ChIP-Seq processing and analyses

After initial quality checks of newly generated data using fastqc, paired-end reads were aligned to the dm6 version of the *Drosophila* genome using bowtie 2 ^7^ (v2.3.5.1) with the following parameters: --local --very-sensitive-local --no-unal --no-mixed --no-discordant --phred33 -I 10 -X 700, and low mapping quality were discarded using samtools ^8^ (-q 30, v1.10, using htslib 1.10.2-3).

PH, H3K27me3, H3K27Ac and H2AK118Ub peaks/domains were called for each replicate separately and on merged reads – using IgG as input control – using macs2 ^9^ (v2.2.7.1) with the following parameters: --keep-dup 1 -g dm -f BAMPE -B --SPMR. Only the peaks that were detected in both replicates (enrichment>0 AND qvalue<0.05) and using merged reads (enrichment>2 AND qvalue<0.01) were retained for further analyses, after being merged with a minimum gap size of 250 bp for narrow peaks (PH and H3K27Ac) and 2.5kb for broad marks (H3K27me3 and H2AK118Ub). macs2 bedgraph files were used for visualization purposes.

PH, H3K27me3 and H2AK118Ub bound genes were defined as overlapping with at least one confident peak (see previous section) anywhere in the body of the gene and up to 2.5kb upstream of its TSS.

For differential analysis of H3K27me3, H3K27Ac and H2AK118Ub, peaks were merge across all conditions (maximum gap of 250 bp for H3K27Ac, 2.5kb for H3K27me3 and H2AK118Ub), overlapping reads were counted using the featureCounts function from the Rsubread R package (v2.0.1, isPairedEnd = TRUE) and differential analysis was performed using the DESeq2 R package (v1.26.0, size factors= total number of aligned reads, design= ∼replicate+condition).

### GO enrichment

Gene ontologies were carried out using the AnnnotationDbi R packge (https://bioconductor.org/packages/AnnotationDbi, v1.48.0). Briefly, genes of interest were compared to a universe set of genes consisting of all genes that passed DESeq2 initial filters, using one-sided Fisher’s exact test (alternative = “greater”). Obtained p-values were corrected for multiple testing using Benjamini-Hochberg’s method.

### Motif enrichment

Non-redundant TF motifs were obtained from de Almeida et al. ^10^, and only the motifs that are associated to a known Drosophila TF were considered. Motif were counted at the promoters of reversible and irreversible genes as well as unaffected genes (TSS -200/+50bp) using the matchMotifs function from the motifmatch R package (Schep A (2022). motifmatchr: Fast Motif Matching in R. R package version 1.18.0) with the following parameters: p.cutoff = 5e-4, bg = “genome”, genome= “dm6”. Then, both sets were compared using two-sided Fisher’s exact test and p-values were corrected for multiple testing using Benjamini-Hochberg’s method. In the case where a single TF would be associated to several motifs, only the most significant value was reported.

### Analysis of Human solid tumors

The differential gene expression analysis was carried out by employing Mann-Whitney test using the TNMplot database which contains transcriptome-level RNA-seq data for different tumor samples from The Cancer Genome Atlas (TCGA) and The Genotype-Tissue Expression (GTEx) repositories (https://pubmed.ncbi.nlm.nih.gov/33807717/).

The survival analysis was carried out using the Pan-Cancer (Bladder, Lung adenocarcinoma, and Rectum adenocarcinoma) or gene-array (Breast, Ovarian, and Prostate) (https://pubmed.ncbi.nlm.nih.gov/34527184/ and https://pubmed.ncbi.nlm.nih.gov/34309564/) datasets of the online tool www.kmplot.com (accessed on 22 December 2022). The Pan-Cancer dataset is based on TCGA data generated using the Illumina HiSeq 2000 platform with survival information derived from the published sources (https://pubmed.ncbi.nlm.nih.gov/33723286/). The gene-array samples were obtained using Affymetrix HGU133A and HGU133plus2 gene chips. The samples were MAS5 normalized and the mean expression in each sample was scaled to 1000. The most reliable probe sets to represent single genes were identified using JetSet (https://dash.harvard.edu/handle/1/10295333).

In the survival analysis, each cutoff value between the lower and upper quartiles of expression was analyzed by Cox proportional hazards regression and false-discovery rate was computed to correct for multiple hypothesis testing. Then, the best performing cutoff was used in the final analysis. Kaplan-Meier survival plots were generated to visualize the survival differences, and hazard rates with 95% confidence intervals were computed to numerically assess the difference between the two cohorts. The statistical analysis was performed in the R statistical environment (www.r-project.org). The analysis results for single genes can be validated using the platforms at www.kmplot.com and www.tnmplot.com.

### Analysis of Multiple Myeloma patient cohorts

For gene expression profiling data from patients with Multiple Myeloma, we used six cohorts that included Affymetrix gene expression data (HGU133plus2) of purified MM cells from the TT2 (Gene Expression Omnibus, accession number GSE2658), TT3 (accession number E-TABM-1138 accession number GSE4583) and Hovon (accession number GSE19784) cohorts (345, 158 and 282 newly-diagnosed MM patients treated by high-dose melphalan and autologous hematopoietic stem cell transplantation); the Mulligan cohort (188 patients at relapse treated by proteasome inhibitor in monotherapy); the Mtp cohort non eligible to HDT ^11^ (63 newly-diagnosed MM patients non eligible to high-dose melphalan and autologous hematopoietic stem cell transplantation) and the Mtp Dara cohort ^11^ (51 patients at relapse treated by anti-CD38 monoclonal antibody (Daratumumab)). Gene expression data were normalized with the MAS5 algorithm and processing of the data was performed using the webtool genomicscape (http://www.genomicscape.com), as done previously ^12,13^, using the R environment (www.r-project.org). The prognostic value of PHC1, PHC2, PHC3, CBX2, CBX7 et BMI1 gene expression was investigated using the Maxstat R function and Kaplan Meier survival curves as previously described ^14^. The differential gene expression analysis between normal bone marrow plasma cells from healthy donors and MM cells from patients was carried out by employing Mann-Whitney test. The prognostic value of *PHC1, PHC2, PHC3, CBX2, CBX7* and *BMI1* genes was combined using our previously published methodology ^14^ (sum of the Cox b-coefficients of each of the 6 genes, weighted by ±1 if the patient MMC signal for a given gene is above or below the probe set Maxstat value of the gene). Clustering was performed using the Morpheus software (https://software.broadinstitute.org/morpheus) and violin plots using GraphPad Prism software (http://www.graphpad.com/scientific-software/prism/).

## Acknowledgements

We thank Montpellier Resources Imagerie facility as well as the *Drosophila* facilty (both affiliated to BioCampus University of Montpellier, CNRS, INSERM, Montpellier, France). V.P. was supported by the EpiGenMed cluster of Excellence funding (PIA of the French Ministry of Higher Education and Research) and by la Ligue Nationale Contre le Cancer. V.L. was supported by the EpiGenMed cluster of Excellence funding (PIA of the French Ministry of Higher Education and Research). A-M.M was supported by a grant of the Fondation ARC (contract N. 216574, acronym “Epicancer). B.S.. was supported by INSERM. G.C. was supported by CNRS. Research in the G.C. laboratory was supported by grants from the European Research Council (Advanced Grant 3DEpi), the European CHROMDESIGN ITN project (Marie Skłodowska-Curie grant agreement No 813327), the European E-RARE NEURO DISEASES grant “IMPACT”, by the Agence Nationale de la Recherche (PLASMADIFF3D, grant N. ANR-18-CE15-0010), by the Fondation pour la Recherche Médicale (DEI20151234396), by the MSD Avenir Foundation ((Project GENE-IGH), and by the French National Cancer Institute (INCa, PIT-MM grant N. INCA-PLBIO18-362).

## Author contributions

V.L, V.P., A-M.M., and G.C. initiated and led the project. V.P, L.F. and V.L. performed genetic experiments. V.P. performed immunostaining, molecular biology and genomic experiments. V.L. and M.DS. performed computational analysis of genomic datasets. B.S. performed ChIP-seq and CUT&RUN experiments. M.E., B.G and D.C. performed computational analysis of different tumor types. J.M. performed computational analysis of Multiple Myeloma samples. V.L, V.P., A-M.M and G.C. wrote the manuscript. All the authors discussed the data and reviewed the manuscript.

## Data availability

The datasets generated during and/or analyzed during the current study will be available from GEO.

## Supplementary information

This manuscript includes supplementary information, namely ten Extended Data Figures and three Extended Data Tables.

Extended Data Table 1: Differential analyses and FPKMs of no ph-KD, transient ph-KD and constant ph-KD in the fly line expressing GFP marker only during depletion of PH.

Extended Data Table 2: Clustering of differentially expressed genes and recovery status.

Extended Data Table 3 : Differential analyses and FPKMs of no ph-KD, transient ph-KD and constant ph-KD in the fly line expressing constitutive GFP

**Extended Data Fig. 1:**
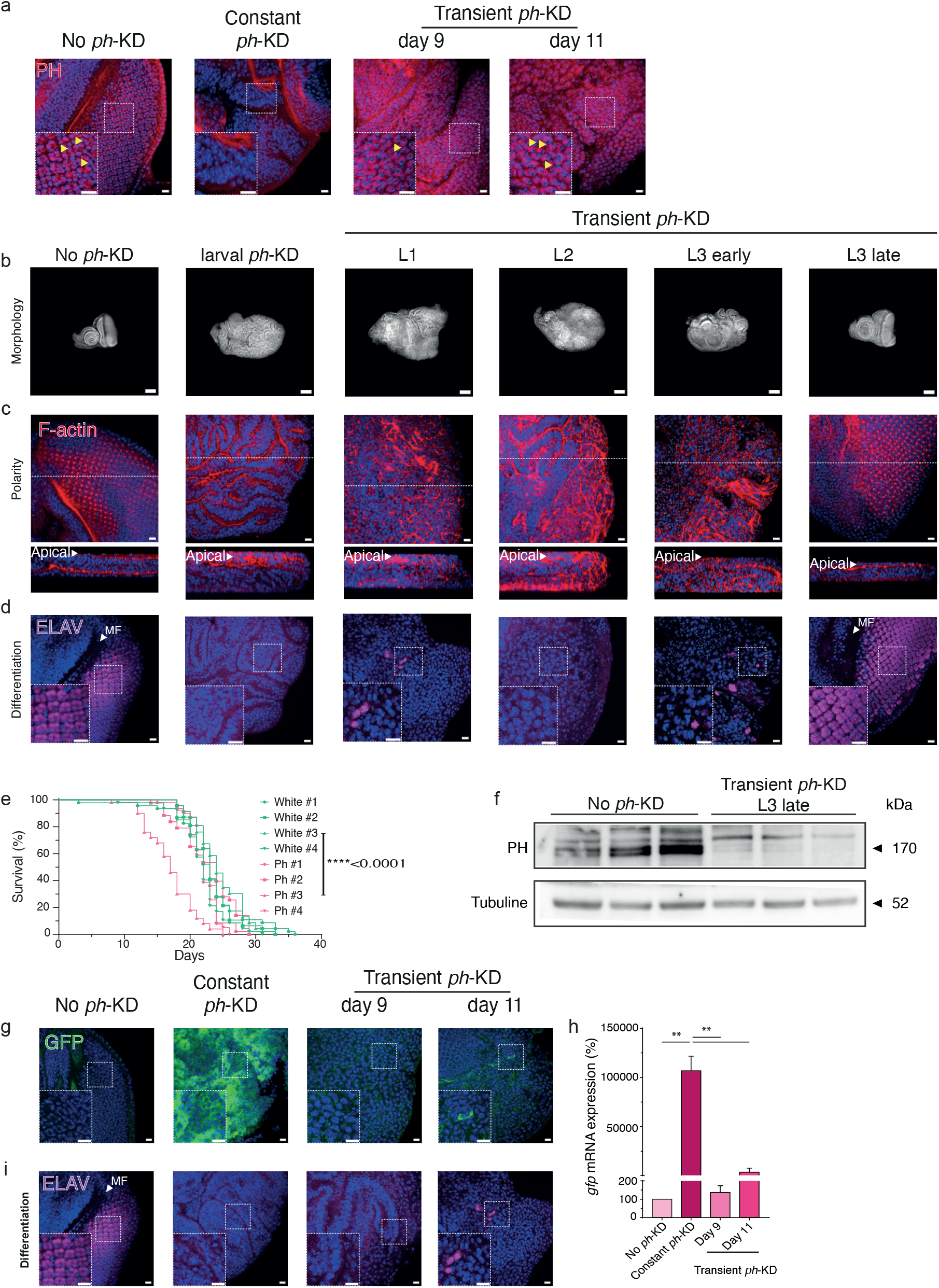
Transient *ph*-KD triggers PH depletion and generates neoplastic tumors that persist after PH recovery. **a-** PH immunostaining (in red) after no *ph*-KD (control), constant or transient *ph*-KD. Tissues were counterstained with DAPI (in blue). **b-d** DAPI staining (**b**); F-actin labeling showing apico-basal polarity (in red; **c**) and neuronal marker ELAV (differentiation, in purple, **d**) staining of EDs after no *ph*-KD (control), or a transient 24H *ph*-KD during first (L1), second (L2), early third or late third (L3) larval instar respectively. **e-** Survival curves of no-*w*-KD and no-*ph*-KD flies, maintained at 29°C only at the adult stage. **f-** Western blot showing PH protein from EDs subjected to no *ph*-KD (control) or transient *ph*-KD at late third instar larval stage in biological triplicate. **g-** GFP staining (in green) after no *ph*-KD (control), constant or transient *ph*-KD. Tissues were counterstained with DAPI (in blue). **h-** RT-qPCR experiments showing *gfp* expression level after no *ph*-KD (control), constant or transient *ph*-KD. Error bars represent the standard error of the mean (SEM) for three independent experiments. ** <0.01. **i-** ELAV (differentiation, in purple) staining after no *ph*-KD (control), constant or transient *ph*-KD. Tissues were counterstained with DAPI (in blue). Scale bars: 10 μm (a, c, d, g, i), 100 μm (b).

**Extended Data Fig. 2:**
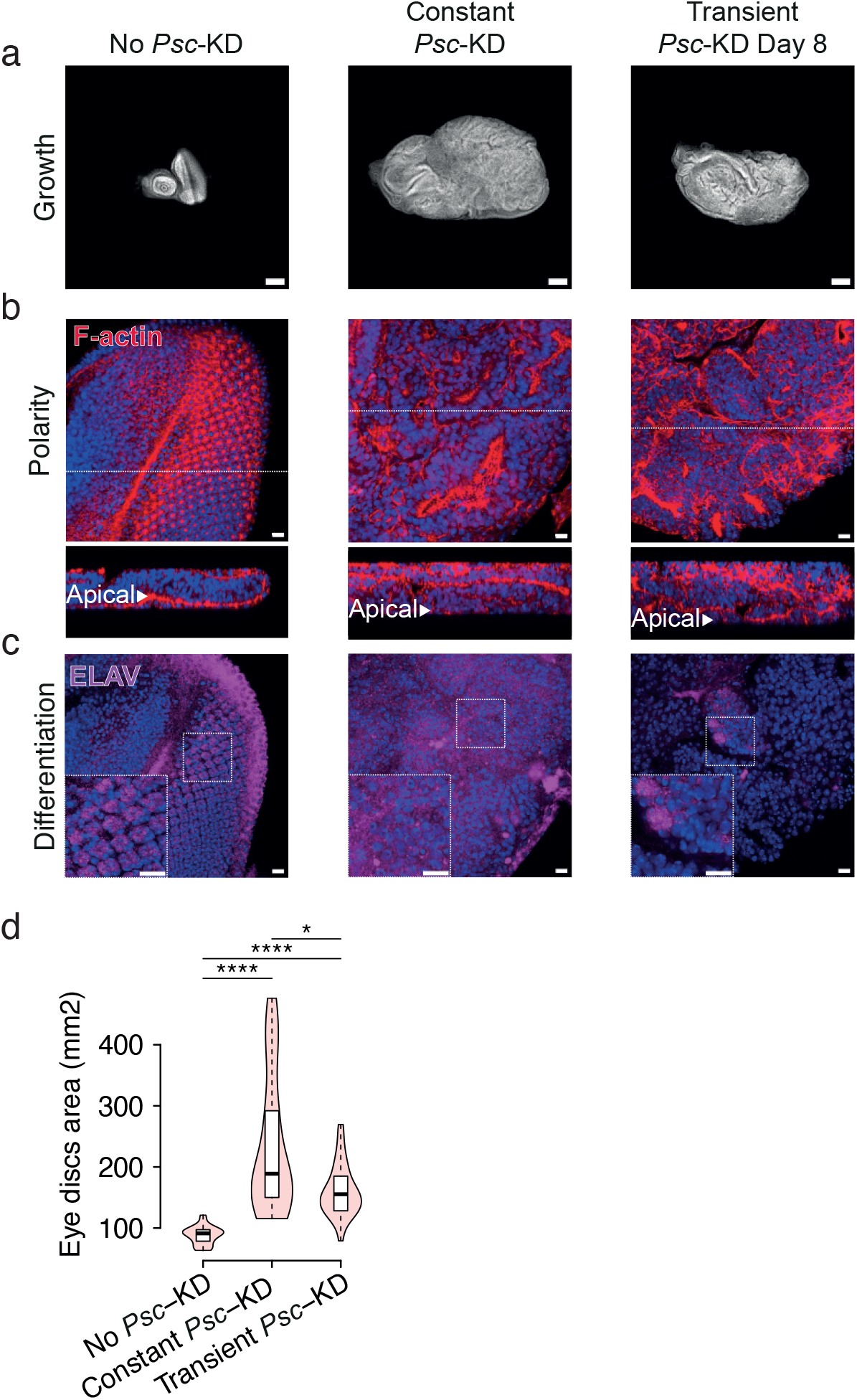
Transient *Psc*-KD phenocopies transient *ph*-KD. **a-c** DNA is stained with DAPI (blue) in EDs under the indicated conditions (**a**); F-actin staining (in red; **b**) showing apico-basal polarity and ELAV neuronal marker (differentiation, in purple, **c**) staining after no *Psc*-KD, constant or transient *Psc*-KD. **d-** Violin plots showing a comparative measure of ED size quantified as an overall DAPI staining area under no *Psc*-KD (control), constant or transient *Psc*-KD conditions (Wilcoxon test, *pval<5e-2, ****pval<1e-5).

**Extended Data Fig. 3:**
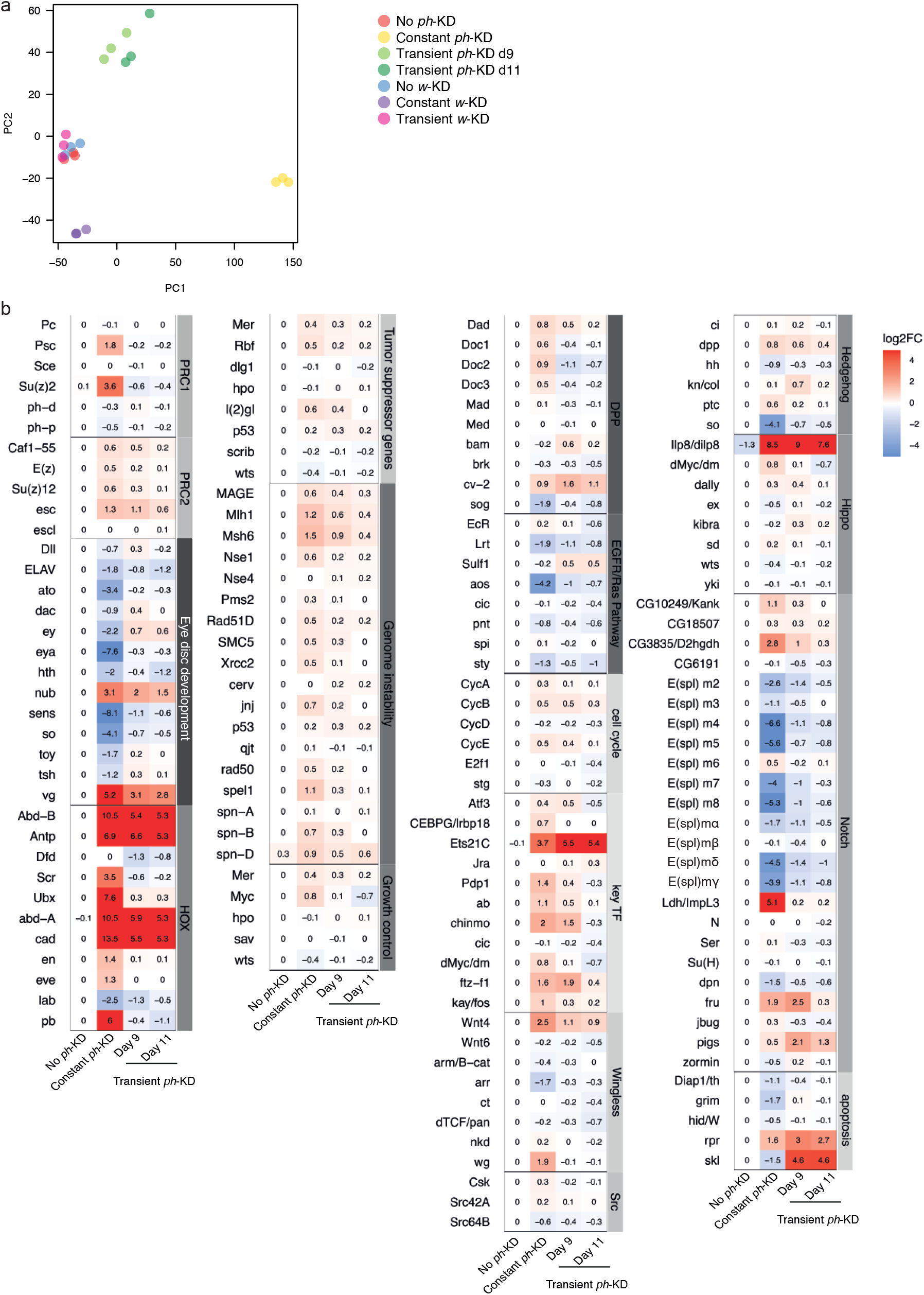
Transcriptomic analysis of the effect of a constant or transient *ph*-KD. **a-** Principal Component Analysis (PCA) on the biological triplicate of gene expression data. Each dot represents one sample. PCA shows the high similarity between controls no *w*-KD and no *ph*-KD. **b-** Trancriptional fold changes upon PH depletion of genes known to be involved in tumorigenesis in Drosophila: PcG members, differentiation and Hox genes, tumor suppressor genes, genes involved in genome instability, cell cycle, apoptosis and growth control, members of major signaling pathways (Dpp, Decapentaplegic; EGFR, Epidermal Growth Factor Receptor; Notch; Hh, Hedgehog; Wg, Wingless; Hippo).

**Extended Data Fig. 4:**
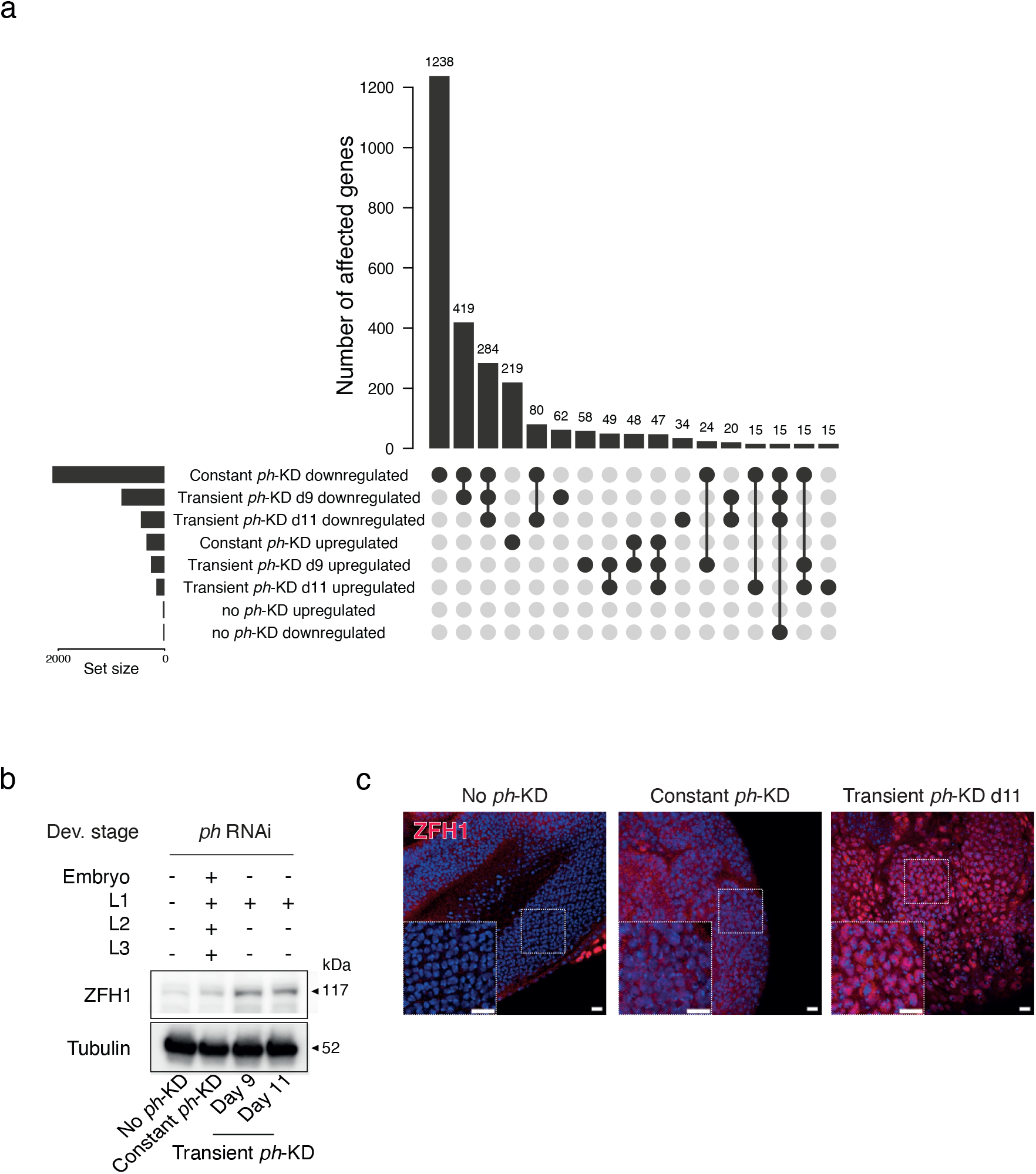
ZFH1 is included among irreversibly derepressed genes after transient PH depletion. **a-** Representation of the number of overlaps for transcriptionally down-regulated or up-regulated genes between the indicated conditions. **b-** Western blot showing ZFH1 in EDs subjected to no *ph*-KD (control), constant or transient *ph*-KD. **c-** ZFH1 immunostaining (in red) after no *ph*-KD (control), constant or transient *ph*-KD. Tissues were counterstained with DAPI (in blue). Scale bars: 10 μm.

**Extended Data Fig. 5:**
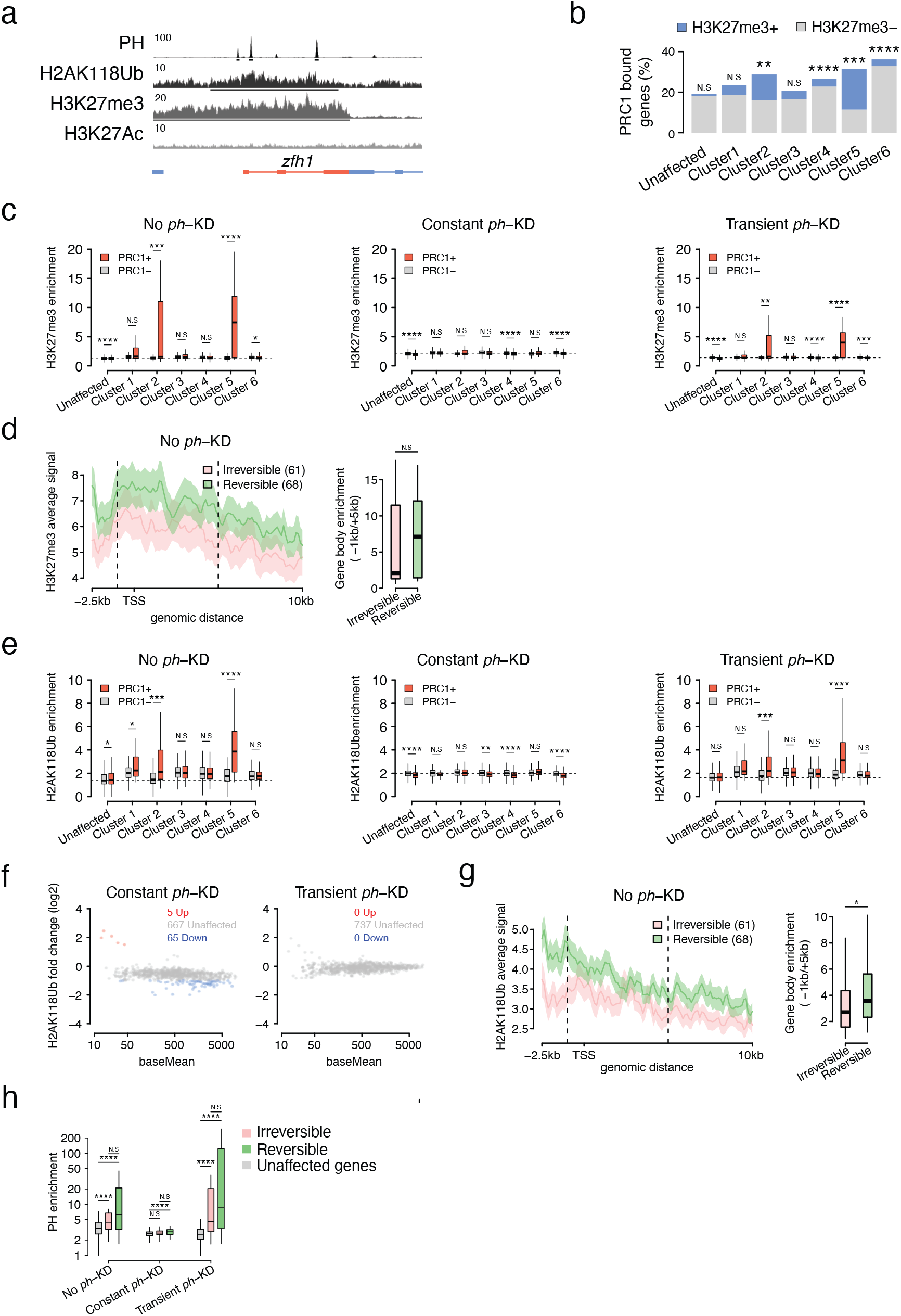
Similar recovery of chromatin landscape at reversible and irreversible genes. **a-** Screenshot of PH ChIP-Seq, H2AK118Ub, H3K27me3 and H3K27Ac CUT&RUNs at the *zfh1* locus in no *ph*-KD (control) EDs. **b-** Proportion of PRC1-bound genes per cluster, using all unaffected genes as a reference. Overrepresentation of PRC1-bound genes was assessed using two tailed Fisher’s exact test (N.S-not significant, *pval<5e-2, **pval<1e-2, ***pval<1e-3, ****pval<1e-5). **c-** H3K27me3 quantification at gene bodies per cluster, further stratified for the presence or absence of PRC1 binding (in orange and grey, respectively). Wilcoxon test, N.S-not significant, *pval<5e-2, **pval<1e-2, ***pval<1e-3, ****pval<1e-5. **d-** Average signal and quantification of H3K27me3 at the body of irreversible (pink) and reversible (green) genes (Wilcoxon test, N.S-Not significant). **e-** H2AK118Ub quantification at gene bodies per cluster, further stratified for the presence or absence of PRC1 binding (in orange and grey, respectively). Wilcoxon test, N.S-not significant, *pval<5e-2, **pval<1e-2, ***pval<1e-3, ****pval<1e-5. **f-** Differential analysis of H2AK118Ub domains that are unaffected (grey) or exhibiting decreased (blue) or increased (orange) enrichment upon constant (left) or transient (right) PH depletion. **g-** Average signal and quantification of H2AK118Ub at the body of irreversible (pink) and reversible (green) genes (Wilcoxon test, N.S-Not significant). **h-** Quantification of PH enrichment at the promoter of irreversible (pink) and reversible (green) genes (Wilcoxon test, N.S-not significant, *pval<5e-2, **pval<1e-2, ***pval<1e-3, ****pval<1e-5).

**Extended Data Fig. 6:**
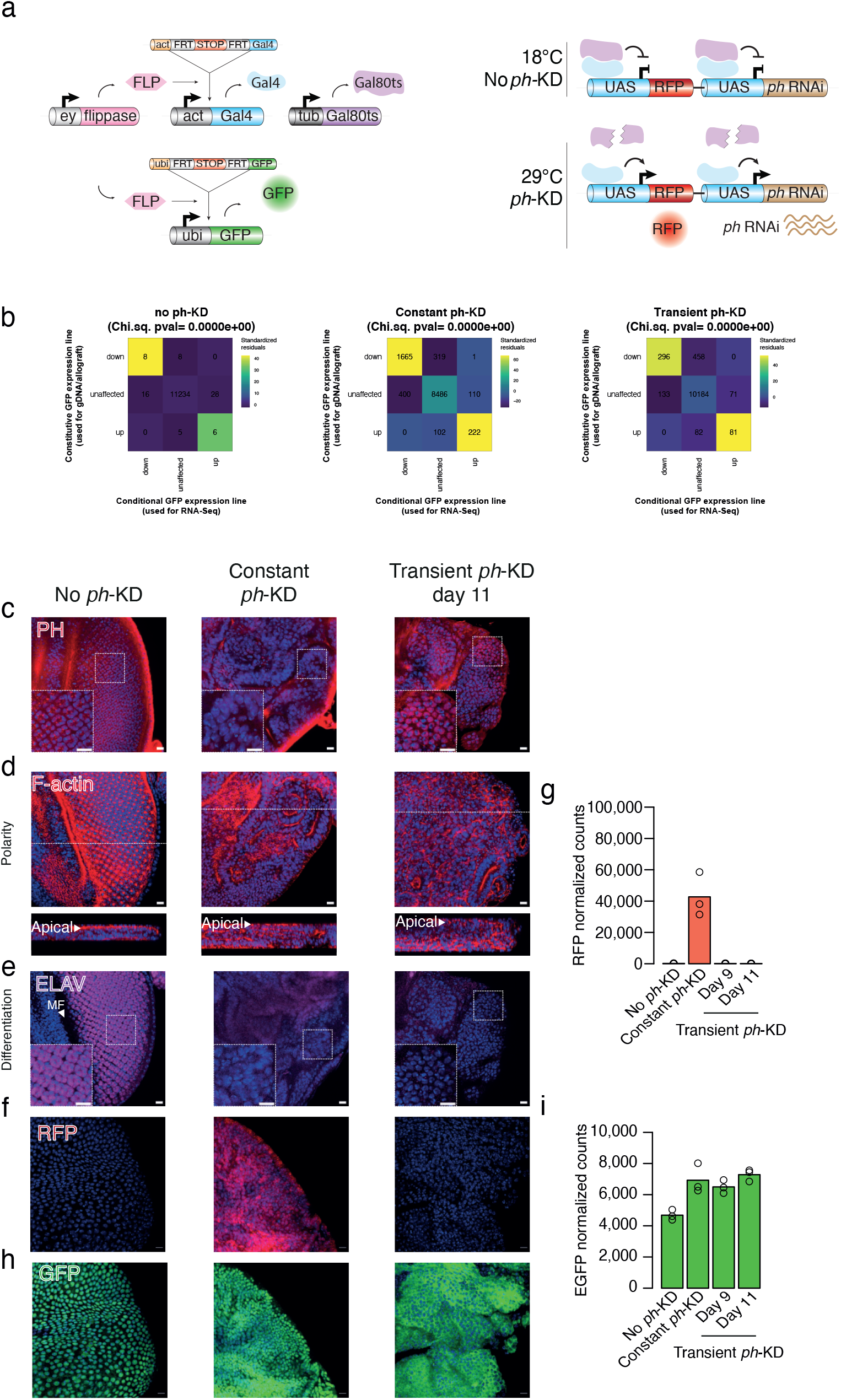
A conditional genetic tool allowing the long-term tracking of cells submitted to constant or transient PH depletion. **a-** Schematic overview of the thermosensitive *ph*-RNAi genetic system used in allograft experiments. This fly line contains two fluorochromes: GFP which is ubiquitously expressed (Ubip63 promoter), in order to follow the tumor evolution through multiple transplantation cycles, while the RFP expression is a read-out of the *ph*-KD which depends, like *ph*-RNAi, on the binding of the yeast transactivator Gal4 to its UAS target sequences. At 18°C, neither *ph*-RNAi nor RFP are expressed because the Gal4 inhibitor Gal80ts is active. At 29°C, the thermosensitive Gal80^ts^ is inactivated. Gal4 binds to UAS sequences allowing the expression of *ph*-RNAi and RFP. **b-** Chi-squared residuals showing the overlap between the transcriptomes of the conditional GFP line (used for RNA-Seq, Fig. 2) and the constitutive GFP line (developed for gDNA and allografts, Fig. 1 and 5). **c-** PH immunostaining (in red) after no *ph*-KD (control), constant or transient *ph*-KD. Tissues were counterstained with DAPI (in blue). **d-** F-actin staining (in red) showing apico-basal polarity after no *ph*-KD (control), constant or transient *ph*-KD. Tissues were counterstained with DAPI (in blue). **e-** ELAV staining (in purple) showing neuronal differentiation after no *ph*-KD (control), constant or transient *ph*-KD. Tissues were counterstained with DAPI (in blue). **f-** RFP immunostaining (in red) after no *ph*-KD (control), constant or transient *ph*-KD makes it possible to track the dynamics of *ph*-KD. Tissues were counterstained with DAPI (in blue). **g-** Normalized read counts of RFP mRNAs after no *ph*-KD (control), constant or transient *ph*-KD showing that transcriptional expression of RFP occurs at constant 29°exposure but is reverted after a transient *ph*-KD. **h-** GFP staining (in green) after no *ph*-KD (control), constant or transient *ph*-KD. As expected reveal, GFP is expressed in all conditions. Tissues were counterstained with DAPI (in blue). **i-** Normalized GFP read counts after no *ph*-KD (control), constant or transient *ph*-KD showing constant expression of GFP in all conditions. Scale bars: 10 μm.

**Extended Data Fig. 7:**
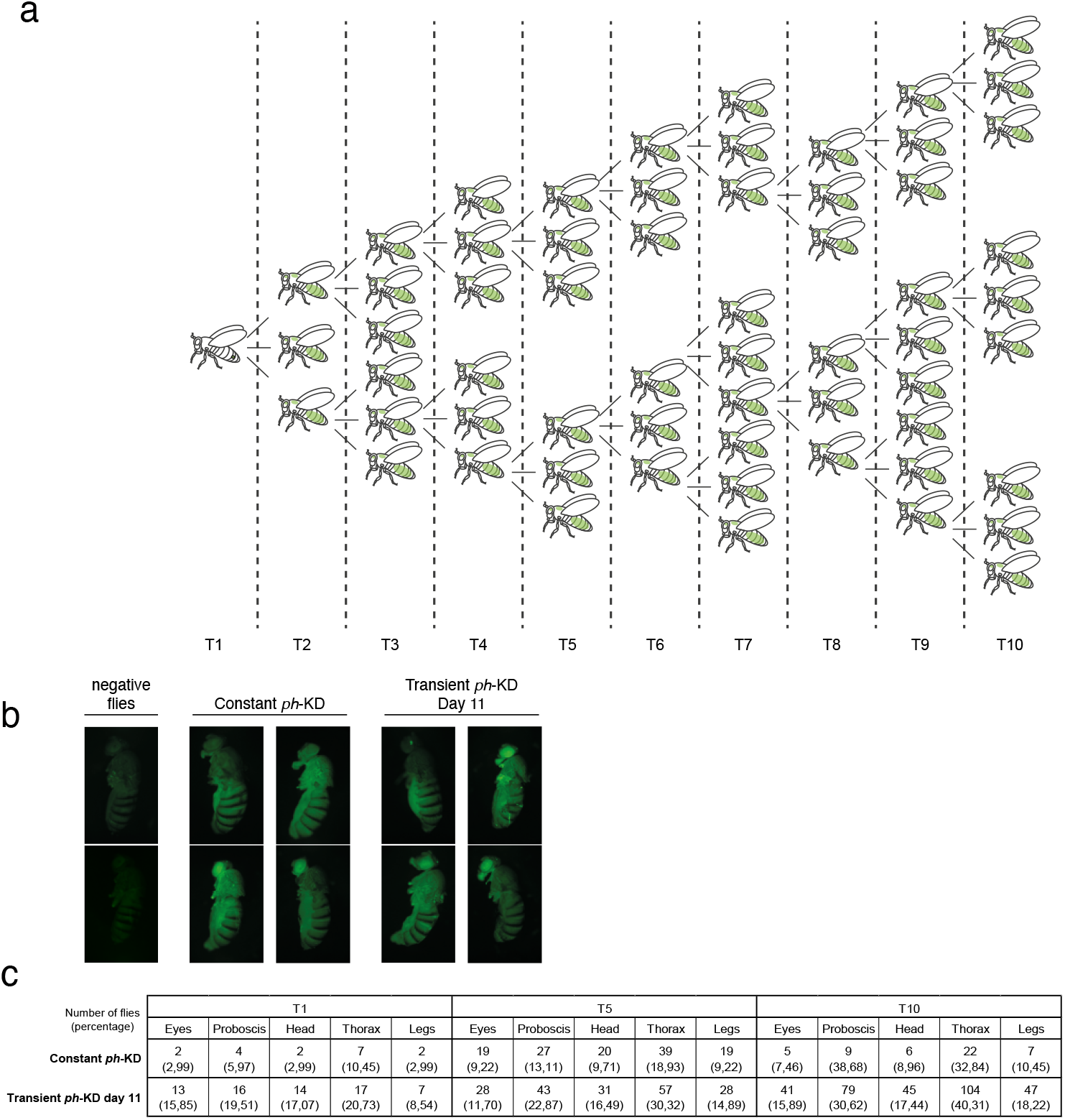
Comparative analysis of metastasis formation through serial transplantation of constantly PH-depleted tumors versus EICs. **a-** Tree representation of the allograft assay. A primary ED tumor derived from a constant or transient *ph*-KD is dissected from donor larvae and repeatedly allografted into the abdomen of a female host maintained at 18°C to prevent re-expression of *ph*-RNAi. Each injected fly is monitored every two days. When the host fly abdomen is completely filled with GFP positive cells, the host is dissected and tumor cells are again injected into multiple hosts. The procedure was repeated up to the T10 generation. **b-** Pictures of flies injected with grafts obtained from no *ph*-KD (control), constant or transient *ph*-KD conditions. The primary tumor can invade the abdomen and surrounding tissues. **c-** In order to score the frequency of metastases, injected flies were monitored twice a week and the appearance of metastasis in the thorax, head, proboscis, eyes and legs were noted for each generation. Values in the table represent the number and percentage (in brackets) of flies with metastases after transplantation of the 1^st^, 5^th^ or 10^th^ (T1, T5 and T10, respectively) generation of constant or transient *ph*-KD tumor transplants.

**Extended Data Fig. 8:**
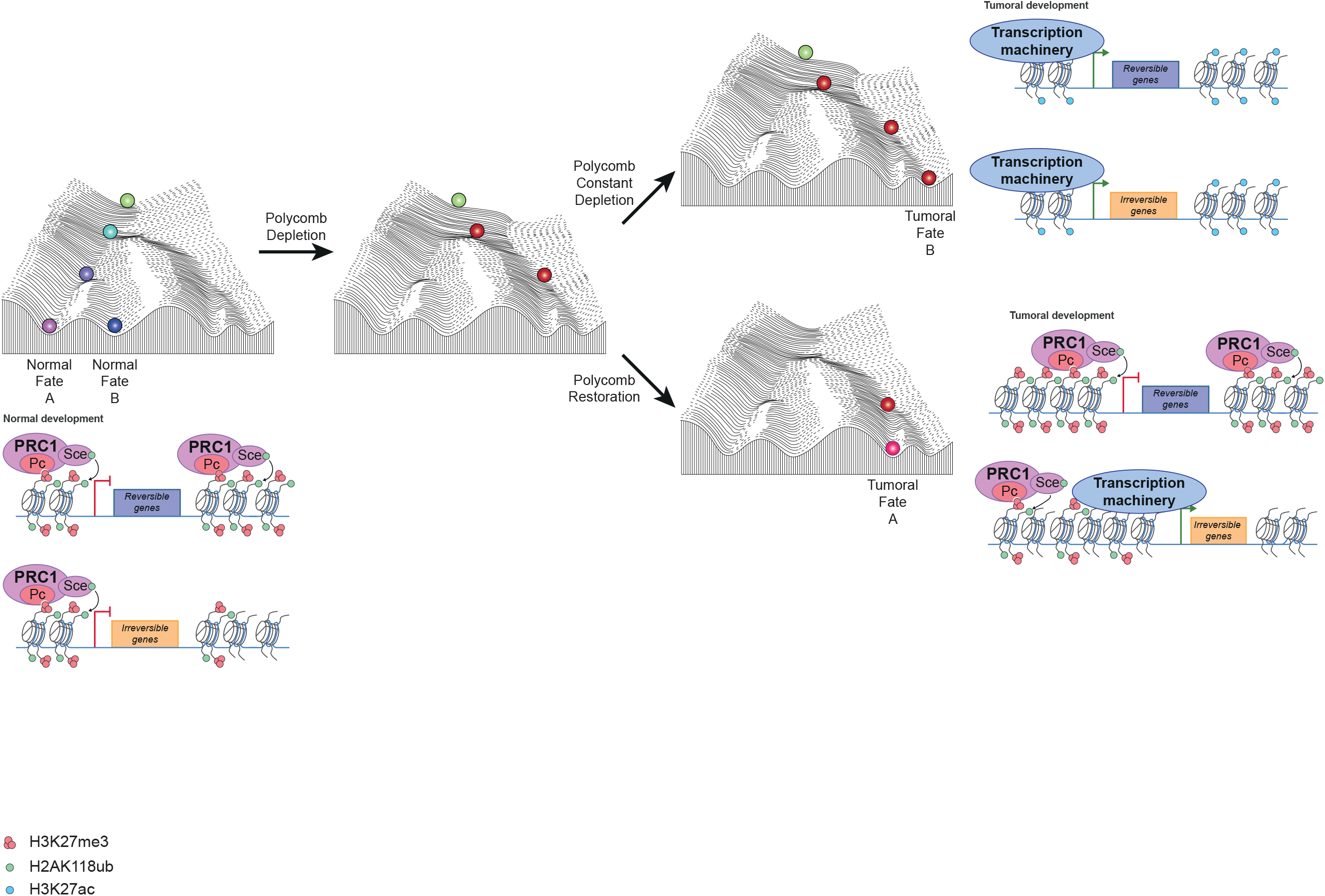
A model explaining the emergence of epigenetically induced cancers. The model is based on the well-known Waddington landscape depicting a marble rolling down a slope with multiple choices of trajectories that depend on the hills and valleys encountered along their path. This scheme is a metaphor of the multiple possible cell fates that can arise from a single cell representing the zygote and is frequently used to signify that epigenetic inheritance contributes to the stable transmission of cell fates, once they are determined by intrinsic and extrinsic signals. In the context of this work, we posit that Polycomb components contribute to shaping the landscape and allow for multiple normal cell fates to be set up and transmitted through the developmental process. In normal development, cells (in green) on top of the hill, will go down the hill during differentiation in order to acquire normal fates. (left panel). Upon depletion of a Polycomb component, such as the PRC1 subunits PH or PSC, the landscape is modified (center panel). If depletion is stably maintained, the modified landscape forces cells to enter a pathway that is both aberrant and intrinsically stable, inducing cancer formation through loss of cell differentiation, loss of cell polarity and sustained proliferation (top right panel). If Polycomb protein levels are restored, the landscape goes back to its original shape. However, if restoration of the landscape occurs after the cells have already chosen an aberrant route (represented by the marble in the middle of the landscape), they can no longer find the healthy trajectory and will be obliged to choose among a limited set of possibilities within a diseased cell space. This ultimately can lead to the maintenance of tumoral phenotypes. In additional to the Waddington landscape panels, gene panels are added along the panels, representing a putative molecular explanation of the phenomenon described here. The chromatin and functional state of reversible and irreversible genes is shown in each condition. In the absence of Polycomb depletion, reversible genes are characterized by the presence of high levels of Polycomb (only PRC1 complexes are shown for simplicity) and by low levels or lack of transcription. Irreversible genes are also not or lowly expressed, but they contain less Polycomb in their chromatin. Removal of PRC1 subunits induces chromatin opening and derepression of both classes of genes. Upon restoration of Polycomb components however, a difference emerges. High levels of Polycomb binding might restore a silent chromatin state, whereas lower levels of Polycomb binding might be unable to restore repression at irreversible genes.

**Extended Data Fig. 9:**
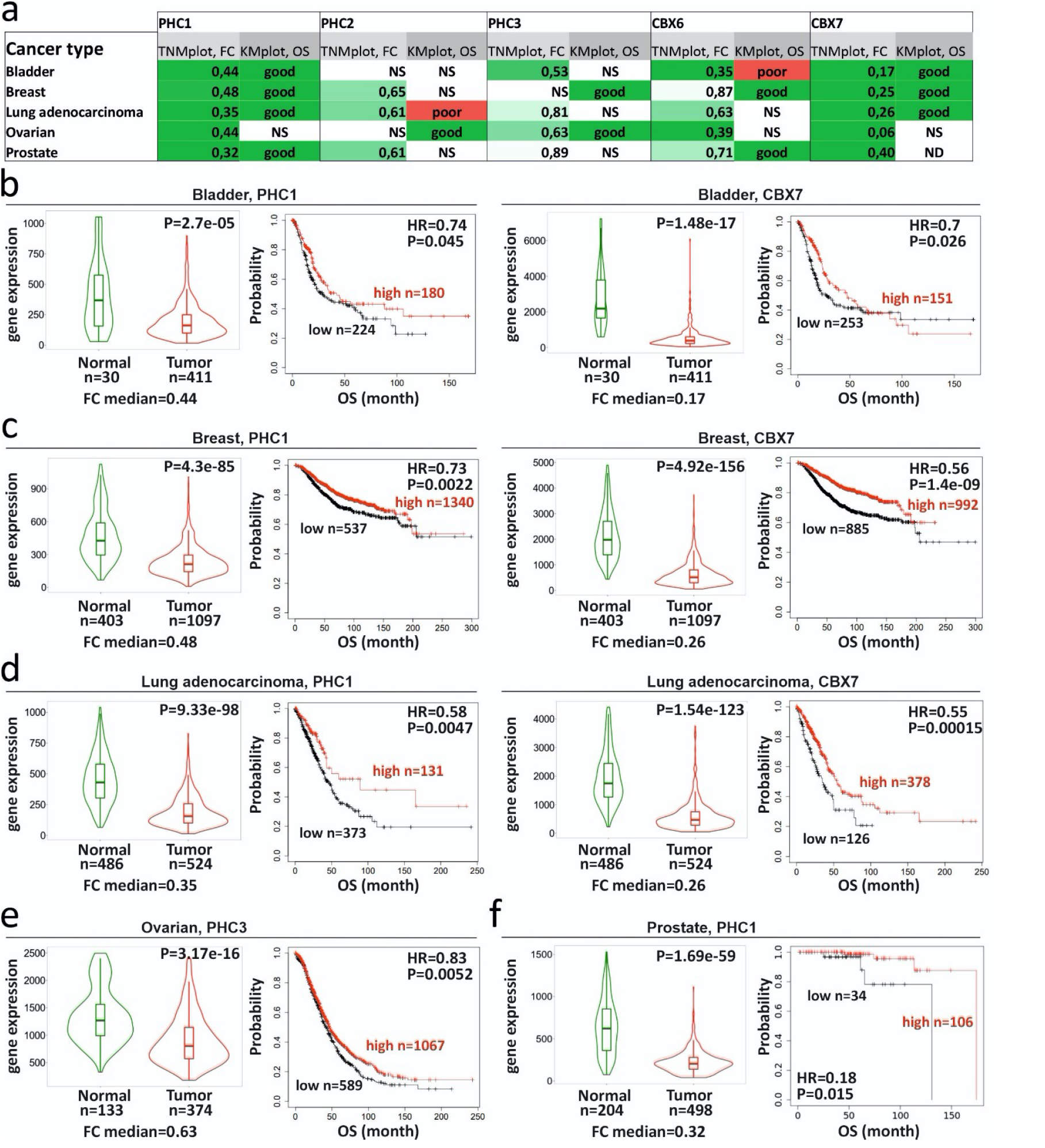
Examples of tumor suppressive role of canonical PRC1 core subunits in different cancer types. **a-** Clinical correlations for PRC1 in selected cancer types. Differential gene expression (TNMplot) and clinical prognosis Kaplan-Meier plot (KMplot) results are given for *PHC1, PHC2, PHC3, CBX6, CBX7* genes. TNMplot columns represent the differential gene expression analysis in tumor and matched normal tissues, which was performed using the https://tnmplot.com/ online tool. FC median: Fold change median. Statistical significance was calculated by a Mann–Whitney U test and set at 0.01. NS – non-significant Mann–Whitney p-value. Green boxes indicate that gene expression is significantly lower in tumor tissues. KMplot columns show the analysis of correlation between the overall survival (OS) and the levels of gene expression. KMplot analysis was performed using the www.kmplot.com online tool. HR: hazard ratio. Statistical significance was calculated by a logrank p-value and was set at 0.05. NS – non-significant logrank *p-*values. Green boxes (“Good”) indicate cases in which high expression of PRC1 genes in tumors is associated with a better overall patient survival. **b-** Clinical prognosis for PHC1 (left) and CBX7 (right) in bladder cancer. For each gene the TNMplot (Violin plots, left panels) and KM plots (right panels) are shown. **c-** Clinical prognosis for PHC1 (left) and CBX7 (right) in breast cancer. For each gene the TNMplot (Violin plots, left panels) and KM plots (right panels) are shown. **d-** Clinical prognosis for PHC1 (left) and CBX7 (right) in lung adenocarcinoma. For each gene the TNMplot (Violin plots, left panels) and KM plots (right panels) are shown. **e-** Clinical prognosis for PHC3 (left) in ovarian and PHC1 (right) in prostate cancer. For each gene the TNMplot (Violin plots, left panels) and KM plots (right panels) are shown.

**Extended Data Fig. 10:**
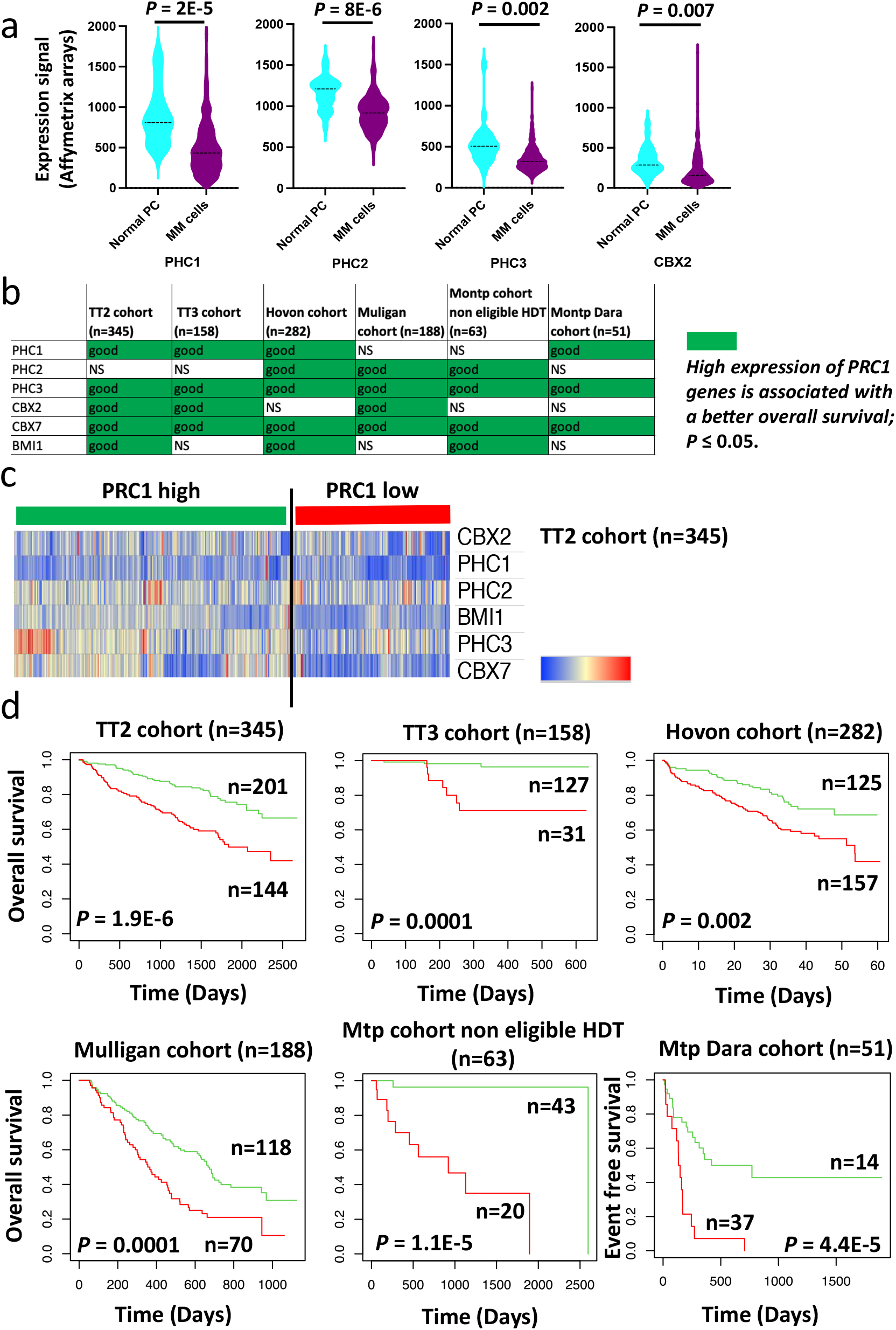
A tumor suppressive role of PRC1 core subunits in Multiple Myeloma. **a-** *PHC1, PHC2, PHC3* and *CBX2* gene expression is significantly downregulated in malignant plasma cells (PCs) from patients with Multiple Myeloma (MM) compared to normal bone marrow PCs. Affymetrix U133 P gene expression profiles of purified bone marrow PC from 22 healthy donors and purified myeloma PCs from 345 previously untreated patients were compared using publicly available data (Gene Expression Omnibus, accession number GSE2658) from the University of Arkansas for Medical Sciences (UAMS, Little Rock, AR). Statistical difference was assayed using a Student t test. **b-** Prognostic value of the PRCA core components in MM. The prognostic value of PHC1, PHC2, PHC3, CBX2, CBX7 et BMI1 gene expression was analyzed in 6 independent cohorts of patients with MM using the Maxstat R function and Kaplan Meier survival curves as previously described. High expression of *PHC1, PHC2, PHC3, CBX2, CBX7* and *BMI1* was associated with significantly shorter overall survival in at least three independent cohorts of MM patients out of the six investigated (green color). The six cohorts included gene expression data of purified MM cells from the TT2, TT3 (accession number E-TABM-1138) and Hovon (accession number GSE19784) cohorts (345, 158 and 282 newly-diagnosed MM patients treated by high-dose melphalan and autologous hematopoietic stem cell transplantation); the Mulligan cohort (188 patients at relapse treated by proteasome inhibitor in monotherapy); the Mtp cohort non eligible to HDT (63 newly-diagnosed MM patients non eligible to high-dose melphalan and autologous hematopoietic stem cell transplantation) and the Mtp Dara cohort (51 patients at relapse treated by anti-CD38 monoclonal antibody (Daratumumab). **c-**The prognostic information of *PHC1, PHC2, PHC3, CBX2, CBX7* and *BMI1* genes was combined. Patients of the TT2 cohort (n = 345) were ranked according to the increased value of the calculated score and a cluster was defined. **d-** In the TT2 cohort, a maximum difference in overall survival was obtained, using the Maxstat R package, splitting patients into high-risk for 144 patients with the lowest expression of PRC1 genes and low-risk group for the 201 patients with higher PRC1 gene expression. Using the same parameter of the TT2 training cohort, we validated the association between low expression of PRC1 genes and a poor outcome in the five other independent cohorts of patients with MM.

**Extended data Table 1: Differential analyses and FPKMs of no *ph*-KD, transient *ph*-KD and constant *ph*-KD (fly line expressing a GFP marker during depletion of PH)**

For each *ph*-KD/*w*-KD comparison, DESeq2 outputs are on separated sheets. “diff”column specifies whether the genes was consider as unaffected, up-or downregulated according to the criteria (| log2FC | > 1 & padj<0.05). Last shee contains FPKMs for all conditions.

**Extended data Table 2: Clustering of differentially expressed genes and recovery status**

For each gene, its corresponding cluster is shown (Figure 2b, “Cluster”column). PH, H3K27me3 and H2AK118Ub binding are also specified and were used to define “Irreversible” and “reversible” genes (see material and methods, “Recovery” column). Finally, PH binding at the gene promoter is shown and was used to restrict “irreversible” and “reversible” gene sets (see material and methods, “PH bound promoter Recovery” column).

**Extended data Table 3: Differential analyses and FPKMs of no *ph*-KD, transient *ph*-KD and constant *ph*-KD (fly line expressing GFP constitutively)**

For each *ph*-KD/*w*-KD comparison, DESeq2 outputs are on separated sheets. “diff”column specifies whether the genes was consider as unaffected, up-or downregulated according to the criteria (| log2FC | > 1 & padj<0.05). Last shee contains FPKMs for all conditions.

